# A modular RNA interference system for multiplexed gene regulation

**DOI:** 10.1101/2019.12.12.873844

**Authors:** Ari Dwijayanti, Marko Storch, Guy-Bart Stan, Geoff S. Baldwin

## Abstract

The rational design and realisation of simple-to-use genetic control elements that are modular, orthogonal and robust is essential to the construction of predictable and reliable biological systems of increasing complexity. To this effect, we introduce **m**odular **A**rtificial **R**NA **i**nterference (mARi), a rational, modular and extensible design framework that enables robust, portable and multiplexed post-transcriptional regulation of gene expression in *Escherichia coli*. The regulatory function of mARi was characterised in a range of relevant genetic contexts, demonstrating its independence from other genetic control elements and the gene of interest, and providing new insight into the design rules of RNA based regulation in *E. coli*, while a range of cellular contexts also demonstrated it to be independent of growth-phase and strain type. Importantly, the extensibility and orthogonality of mARi enables the simultaneous post-transcriptional regulation of multi-gene systems as both single-gene cassettes and poly-cistronic operons. To facilitate adoption, mARi was designed to be directly integrated into the modular BASIC DNA assembly framework. We anticipate that mARi-based genetic control within an extensible DNA assembly framework will facilitate metabolic engineering, layered genetic control, and advanced genetic circuit applications.

## INTRODUCTION

Synthetic biology aims to make the engineering of biological systems more predictable, efficient and reliable (1–3). With this goal in mind, our ability to predictably combine genetic parts into higher-level functional biological/biochemical systems of increasing complexity is essential to improve the efficiency of the Design-Build-Test-Learn cycle (4, 5). In this work, we focus on expanding the toolbox of modular biomolecular control elements that function at the post-transcriptional level, specifically small non-coding RNAs (sRNAs). The ubiquity of sRNA-based control in critical cellular processes such as homeostasis and adaptation demonstrates their importance for the design and control of biological systems (6–10). The sRNAs act to coordinate and synchronise multiple signals through sequence-specific and transient RNA-RNA interactions (11, 12), typically leading to down-regulation of target gene expression (13–16).

Based on the arrangement of sRNAs and their targets, sRNAs can be grouped into *cis* and *trans*-encoded sRNAs. The *cis*-encoded sRNAs are transcribed from the same loci as their target mRNA, but as a reverse complement, thus making a complementary base-pairing to the mRNA they regulate; in contrast, *trans*-encoded sRNAs are located in a distal site. Despite this, *trans*-encoded sRNAs can have efficient regulation of their mRNA targets (14, 17–19). In bacterial systems, *trans*-encoded sRNAs rather than *cis*-encoded sRNAs have been reported to be especially effective in silencing gene expression due to their longer half-life in the cytoplasm, thought to be due to association with the Hfq chaperone protein (20–23). The *trans*-encoded sRNAs, therefore, have been widely applied as versatile genetically encoded controllers for the design of synthetic biological circuits for metabolic engineering and synthetic biology purposes (7, 24–27).

Previous approaches to implement *trans*-encoded sRNA were exemplified by the use of native sRNAs and their cognate targets i.e. *rpoS, hns, sodB*, and *ompC* (28, 29). By fusing the leader sequences from these natural sRNA systems to a reporter gene (e.g. GFP and mCherry), the performance of the native sRNA system could be quantified from the fluorescence output. Due to its highly composable structure, the sRNA can also be re-programmed to target any RNA sequence by modification of the seed sequence, whilst retaining the native sRNA scaffold (17, 18, 28). We term this new type of sRNA an artificial sRNA system. Several rules and guides have been developed to improve the rational design of a set of novel sRNA systems (17, 27, 30, 31). This strategy has been implemented to create artificial sRNA systems with bespoke designs, where the sRNA is targeted to bind the first 24 bp of the GOIs (25, 32). Artificial sRNAs have also been employed to specifically bind a set of standardised target sequences that are inserted upstream of GOIs (24). However, the requirement to insert specific target sites upstream of each GOI may impede the implementation and adoption of sRNA-based controllers for broader applications.

To leverage the use of sRNAs in genetic circuits constructed via a modular DNA assembly method, we set out to design a universal, modular framework for the facile implementation of *trans*-acting sRNA-based control using verified, re-usable target sequences, namely **m**odular **A**rtificial **R**NA **i**nterference (mARi). BASIC DNA assembly provides a modular, standardised and automatable (33) framework for genetic structure based on orthogonal, computationally designed linkers (34) (Figure 1). We exploited this by targeting the DNA linkers used in the construction of gene expression cassettes and operons. In our mARi design, a modified seed sequence specific to the linker upstream of the target Gene of Interest (GOI) was fused to a native sRNA scaffold containing a host factor-1 (Hfq) binding site (17, 27) (**Fig. 1a, 2a**). The integration of mARi-based regulation into a modular DNA assembly method offers a simple yet powerful strategy for implementing sRNA regulatory systems. Since standardised linkers are used in the assembly process, gene expression can be controlled by expression of an mARi that is cognate to the linker sequence upstream of the target gene (**Fig. 1**). We characterise the post-transcriptional regulation of mARi in the context of various genetic design parameters including transcript ratio, molecular copy number, spatial organisation, growth temperatures and media, host strain, and growth phase, thus expanding our understanding of the design rules for new artificial sRNA-based regulators. Finally, mARi sequence variants were further expanded and their implementation was demonstrated for simultaneous regulation in a multi-gene system with different genetic architectures.

**Figure 1.**
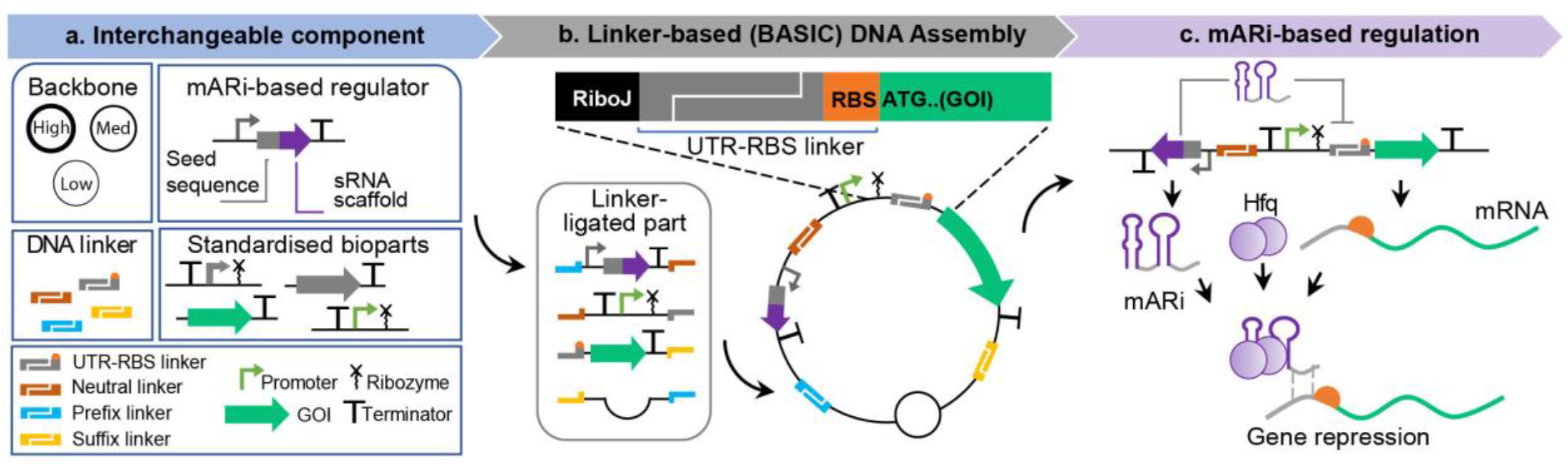
mARi-based regulation integrated into the BASIC modular design and assembly framework. (**a**) mARi-based regulators composed of a promoter, seed sequence (grey), sRNA scaffold (purple) and terminator. These regulators were created as interchangeable components together with standardised DNA parts (i.e. plasmid backbone, promoter, and coding sequence) and computationally designed DNA linkers (i.e. neutral, UTR-RBS, and Prefix/Suffix linker) used in the BASIC DNA assembly framework (34). (**b**) Plasmids were constructed from the interchangeable DNA parts using BASIC DNA assembly: a representative assembled plasmid shows a functional UTR-RBS linker controlling expression of a downstream gene of interest (GOI). (**c**) Expression of mARi leads to repression of a target gene with a cognate UTR by base-pairing between the 5’untranslated region and the mARi seed sequence.

## MATERIALS AND METHODS

### Bacterial strains and outgrowth conditions

The following *E. coli* strains were used for plasmid construction and expression: DH5α (F^−^ *endA1 glnV44 thi-1 recA1 relA1 gyrA96 deoR nupG purB20* ϕ80d*lacZ*ΔM15 Δ (*lacZYA-argF*)U169, hsdR17 (*r* _*K*_ ^−^*m*_*K*_ ^+^), λ^−^), DH10b [F-*mcr*A Δ (*mrr*-*hsd*RMS-*mcr*BC) ϕ80*lac*ZΔM15 Δ*lac*X74 *rec*A1 *end*A1 *ara*D139 Δ (*ara leu*) 7697 *gal*U *gal*K *rps*L *nup*G λ-], BL21 (DE3) (F^−^*omp*T *hsd*S_B_ (r_B_^−^, m_B_^−^) *gal dcm* (DE3)), and BL21star (DE3) (F^−^*omp*T *hsd*S_B_ (r_B_^−^, mB^−^) *gal dcm rne*131 (DE3)). *E. coli* colonies were grown from single-colony isolates in LB (Luria Bertani) medium supplemented with the appropriate antibiotics, shaken at 220 rpm and grown at 37°C (unless specified otherwise). The antibiotics used for plasmid maintenance were carbenicillin and kanamycin at a concentration of 100 μg/ml and 50 μg/ml, respectively.

### Design and analysis of mARi sequences

The sequences of UTR-RBS linkers used in BASIC DNA assembly (34) were designed and validated using R2O DNA designer (35). Secondary structures of the mARi sequences were predicted using RNAFold WebServer (http://rna.tbi.univie.ac.at/cgi-bin/RNAWebSuite/RNAfold.cgi) and visualised using VARNA GUI (36) (http://varna.lri.fr/index.php?lang=en&page=tutorial&css=varna). The predicted interactions between mARi regulators and their mRNA targets were simulated through IntaRNA (37, 38) (https://bio.tools/intarna). The free binding energy of mARi and its mRNA target site was estimated using a web-based service of the Two-State Melting Hybridisation of UNAFold RNA folding software DINAMelt (39) (http://www.unafold.org/Dinamelt/applications/two-state-melting-hybridization.php) using default parameters at 37°C. The percent identity of mARi-UTR pairs was calculated using the EMBOSS needle method (https://www.ebi.ac.uk/Tools/psa/emboss_needle/). The Hamming distance was calculated using a custom Python script (https://github.com/MsDwijayanti/hamming/blob/main/hamming.py). The predicted expression strength of the UTR-RBS linker was simulated using EMOPEC (40) and RBS calculator v2.0 (41, 42) with the input sequence starting from the Transcription Start Site (+1) to 100 bp of the downstream CDS. The off-target effect was computed using CopraRNA (38, 43, 44) (https://bio.tools/coprarna). NC_000913 (*E. coli* MG1655), NC_010473 (*E. coli* DH10b), NC_012892 and NC_012971 (*E. coli* BL21 (DE3)) were used as inputs references for host-background off-target prediction.

### Plasmid assembly

Plasmids were constructed using the BASIC DNA assembly method (34, 45). New DNA parts with prefix and suffix sequences were synthesised as gBlocks by Integrated DNA Technologies (IDT), gene fragments by TWIST Bioscience, or derived by PCR mutagenesis. The gBlocks and gene fragments were directly cloned into pJET1.2 or pUC AmpR_plasmid as BASIC bioparts. PCR mutagenesis for generating new BASIC parts were carried out using Phusion Polymerase (NEB) and phosphorylated oligonucleotides (IDT). Sequence verification of BASIC bioparts was carried out using Sanger sequencing (Source Bioscience). Details of the BASIC bioparts used to construct the plasmids are provided in **Supplementary Table 5**. Linkers for BASIC assembly were provided by Biolegio or IDT (**Supplementary Table 6**). All constructs were transformed into chemically competent *E. coli* DH5α. Maps of plasmids used in this study are provided in **Supplementary Fig. 9**.

### Flow cytometry assay

Cellular expression of sfGFP and mCherry was performed in 96 well plates with biological triplicates for each construct. *E. coli* carrying an empty backbone (containing only an origin of replication and an antibiotic resistance gene) was used as a negative control. *E. coli* colonies were inoculated from glycerol stock into LB media supplemented with appropriate antibiotics. The colonies were grown for 16 h at 30°C and 600 rpm in a benchtop shaker (Mikura). The culture was then diluted 200x (10 µl into 190 µl, then 10 µl into 90 µl) in LB media using an automated liquid handling robot (CyBio Felix). The plate was grown at 37°C and 600 rpm in a benchtop shaker (Mikura). After 6 h of incubation, the cell density was measured by absorption at 600 nm (Abs_600_; not normalised for path length). Abs_600_ typically reached values in the range 0.6–0.7. Two microliters of liquid culture were taken and diluted into 200 µl Phosphate Buffer Saline (PBS) supplemented with 2 mg/ml kanamycin to inhibit further protein synthesis and cell growth. The fluorescence data of single cells for the dual reporters system (**Fig. 4b and d**) was collected using BD Fortessa flow cytometer with a 488 nm excitation laser and filter 530/30 for sfGFP; 561 nm excitation laser and filter 610/20 for mCherry. For the triple reporters system (**Fig. 4c**), fluorescence was measured using Attune NxT flow cytometer with a 488 nm excitation laser and channel BL1 530/30 for sfGFP; 561 nm excitation laser and channel YL2 620/15 for mCherry; 405 nm excitation laser and channel VL1 440/50 for mTagBFP. In total 10,000 events were collected for each sample. The data was stored as an FCS file 3.0 and analysis was done using FlowJo V10. Single cell population gating was performed after plotting FSC-H against SSC-H and histograms of each channel of fluorescence. The chosen gating covered 86-99% of the total population. The fluorescence intensity of the sample was calculated by subtracting the geometric mean fluorescence of the control (strains with empty backbone). The normalised fluorescence was calculated by dividing the background-subtracted mean fluorescence of the sample with mARi by the background-subtracted mean fluorescence of the sample without mARi expression.

### Plate reader assay

For continuous growth assay, overnight liquid cultures (as for flow cytometry assay) were diluted 200x in LB media (10 µl into 190 µl, then 10 µl into 90 µl) using an automated liquid handling robot (CyBio Felix); the plate was then incubated in a microplate reader (Clariostar, BMG Labtech) with continuous shaking at 37°C and 600 rpm for 12-24 h. For different growth temperatures, the DH5α cells with and without mARi expression plasmids were incubated at 30°C or 37°C in LB medium. For different growth media, the engineered DH5α cell were tested in two mediums: LB and EZ MOPS RDM supplemented with 0.2% glucose. For the EZ MOPS RDM media supplemented with 0.2% glucose, 5 µL samples were diluted into 95 µL of fresh media using an automated liquid handling robot (CyBio Felix). The plate was incubated at 37°C and 600 rpm.

Absorbance at 600 nm and GFP fluorescence (F:482-16/F:530-40) were measured every 15 min. The corrected Abs_600_ was calculated by subtracting the mean Abs_600_ of media only control from the Abs_600_ of each well. The fluorescence intensity was corrected by subtracting the mean fluorescence value of negative control at equivalent Abs_600_. The value Fl/Abs_600_ was calculated by dividing the value of corrected fluorescence by corrected Abs_600_. Statistical analysis was calculated in Prism v.8.0 (GraphPad). The values were compared using two-tailed Student’s t-test for unpaired comparisons and one-way ANOVA.

## RESULTS

### Design of a modular Artificial RNA interference regulation system

The mARi sequences were designed with two core components: the seed sequence and an sRNA scaffold containing the Hfq binding site (**Fig. 1a, 2a**). The scaffold sequence is responsible for recognition by the RNA chaperone protein Hfq, which is highly conserved as part of natural cellular regulation (46, 47) in a wide range of organisms (48–51). One of the intensively-studied Hfq-dependent sRNAs, MicC was chosen from sRNA scaffolds naturally found in *E. coli* (52) and *Salmonella enterica* (16). The inclusion of this RNA chaperone binding site in the sRNA structure has previously been shown to one of the best Hfq binding sites to mediate sRNA-based gene repression (30). The nature of the multiple targets of MicC based sRNAs in controlling the expression of outer membrane proteins (OmpC and OmpD) can be exploited to target any mRNA sequence of interest. Based on this architecture, the seed sequences can be modified while preserving the native MicC scaffold (17, 25, 27, 30) (**Fig. 2a**).

**Figure 2.**
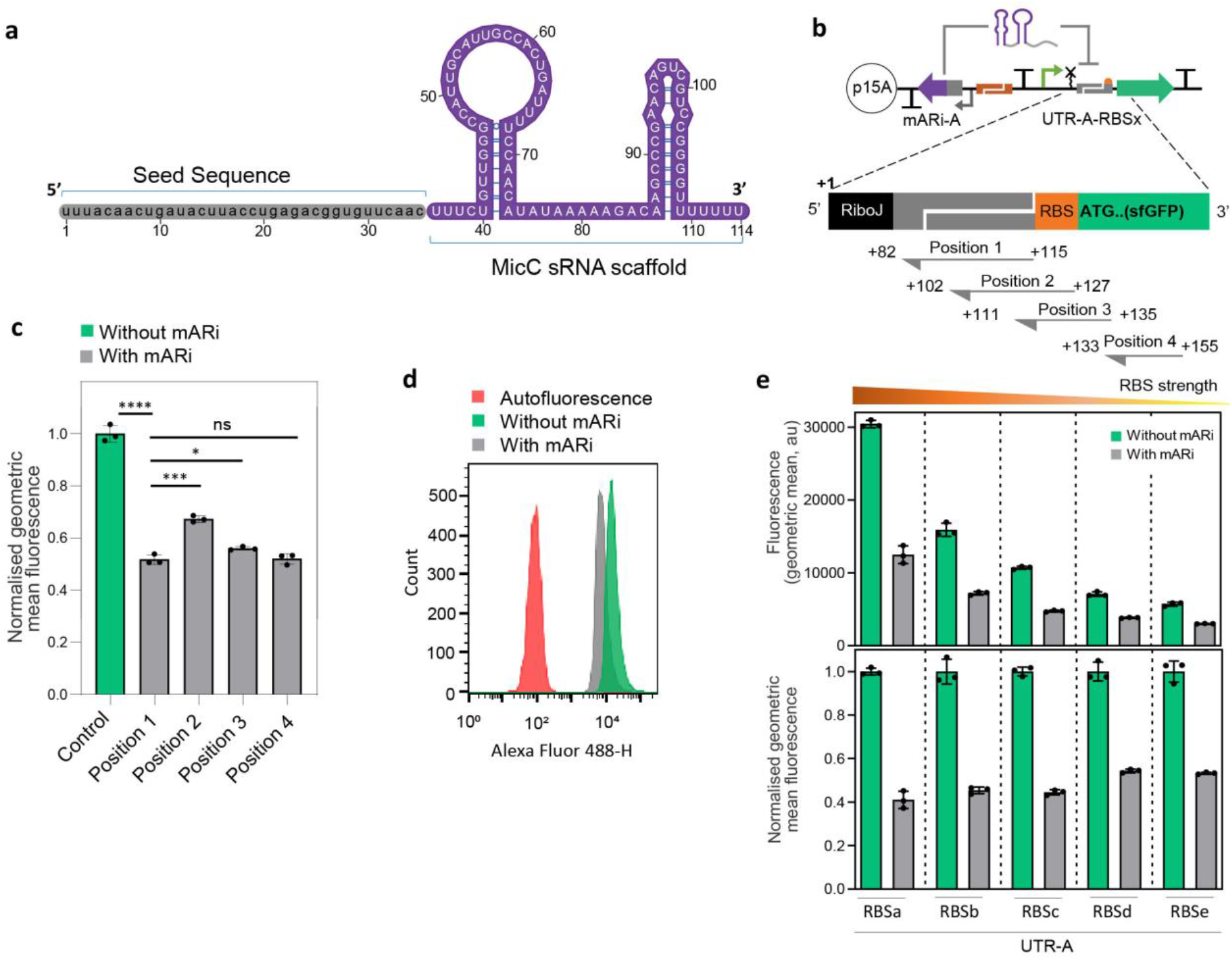
Design and development of mARi-based gene regulation. (**a**) Sequence and secondary structure of mARi containing a seed sequence (grey) and MicC RNA scaffold (purple). (**b**) Schematic design of the target site selection schema for mARi-mediated repression. Four positions within the translation initiation region were selected as target sites (Positions 1-4; **Supplementary Table 1**). The numbers indicate the relative position of mARi downstream of the transcription start site (+1). (**c**) Functional characterisation of mARi-based gene regulation showing the silencing activity of mARi-A for the different target positions vs a sfGFP-only control without mARi. (**d**) Uniformity of representative expressing strains with and without mARi expression. (**e**) The fluorescence intensity (top) and normalized fluorescence (bottom) when position 1 mARi-A was used to target UTR-A with varying RBS strengths. All sfGFP fluorescence measurements were performed by flow cytometry after 6 h growth. Data are shown with error bars for the mean ± SD of triplicate measurements (black dots). Statistically significant differences were determined using two-tailed Student’s t-test (**** represents p<0.0001, *** represents p<0.001, * represents p<0.1, ns represents not significant).

The seed sequence within mARi creates a specific base-pairing interaction to a target sequence in the mRNA of interest, resulting in its post-transcriptional silencing. To standardise and enhance the modularity of mARi, the seed sequence was designed to target expression cassettes constructed via BASIC DNA assembly (34). BASIC assembly relies on standardised linkers between interchangeable DNA parts (**Fig. 1a**) (34). Expression cassettes and operons can be tuned using functional DNA linkers that encode a ribosome binding site (RBS) within a defined 5’ untranslated region (UTR; UTR-RBS linker; **Fig. 1b**). Since BASIC UTRs are computationally designed to be orthogonal to the host genome and exclude secondary structure (35), they provide an ideal target for mARi and reduce off-target effects.

We designed mARi sequences to target the translation initiation region of mRNA for three main reasons: (i) the 5′ UTR sequences are relatively AU rich (30-43.8% GC; **Supplementary Table 1**) and AU rich sequences are preferentially bound by Hfq (53); (ii) targets located in the translation initiation region have been shown to exhibit a low off-target effect in *trans*-encoded sRNA-based regulation (30), which is important for regulator specificity/orthogonality; (iii) a 5′ UTR target site location offers potential modularity since it is independent of different GOIs and is compatible with the modular UTR-RBS linkers in the BASIC design framework (34). To evaluate this approach, seed sequences were designed to target different positions in the translation initiation region (**Fig. 2b**). Positions 1-3 addressed gene-independent targets within the BASIC framework, while position 4 was specific to the gene of interest, in this case *superfolder-gfp* (*sfgfp*; **Fig. 2b**). The reverse complements of the cognate target sequences were combined with the MicC scaffold to create a series of mARi sequences (**Supplementary Table 1**).

The design of the full sequence of each mARi variant was computationally evaluated to ensure that it met the requirements for effective repression (30). Firstly, the designed sequence for each mARi was made compatible with BASIC DNA assembly (34), specifically avoiding the forbidden restriction site *Bsa*I. Secondly, to achieve effective repression and to have a high affinity binding between the seed and target sequences, the GC content and binding energy of the seed sequences were calculated and found to vary from 34.62% to 43.8% and -38.6 kcal/mol to -61 kcal/mol, respectively. The binding energy has a positive correlation to the binding affinity and observed repression capability (27, 30), and previous studies suggested a typical binding energy of -30 to -40 kcal/mol (17, 27) was required for effective repression activity. However, lower binding energy (less than -40 kcal/mol) sRNAs have been experimentally validated and suggested to achieve higher repression (30). Homology length determines mARi specificity and target discrimination (30): shorter binding sites have a lower binding affinity and may cause off-target effects (17, 27), while longer binding sequences may have a complex secondary structure that could cause binding to the mRNA target to be less thermodynamically favorable (30). All mARi regulators used here were designed to have perfect complementarity in their seed sequences to their target mRNA. (**Supplementary Fig. 1b**). To investigate the repression efficiency of mARi targets, sfGFP expression cassettes were assembled with a constitutive promoter P_J23101_BASIC_ and UTR-A-RBSc linker, while mARi expression was driven by the constitutive P_J23101_ (**Fig. 1b**). Measurement of reporter expression demonstrated that all mARi target sequences led to repression of sfGFP, compared to control cells lacking mARi (**Fig. 2c**). The intra-gene target at position 4 showed significant repression as anticipated (17, 25, 27). However, targeting this position would require bespoke design of a specific sequence for the target gene, thus optimisation for each new expression target is required; a less desirable strategy from a modular engineering perspective. Position 1, which targets the UTR upstream of the RBS was as effective as position 4 (**Fig. 2c**). The position 1 target shown had been further optimised for length; reducing the length to 25 bp, as shown for the other positions, had only a minor impact, although further reduction to 20 bp resulted in significant loss of repression activity (**Supplementary Fig. 2**). Targeting this standardised UTR component of the BASIC framework is ideal for modular engineering since it is independent of both the GOI and RBS. The homogeneity of representative producing strains with mARi position 1 against the control strains without mARi expression was shown in **Fig. 2d**.

To evaluate the independence of position 1 UTR target, a set of constructs with five different RBS strengths (RBSa-RBSe) (**Supplementary Table 2**) controlling sfGFP translation were assembled both with and without the position 1 mARi cassette all within the same UTR-A context. The *sfgfp* gene-only constructs resulted in a series of fluorescence intensities reflecting RBS strength variation (**Fig. 2e**). The position 1 UTR-A targeted mARi cassette reduced the fluorescence signal in all cases and the relative repression was found to be constant across the series (**Fig. 2e**). This demonstrated that the repression activity of mARi-A at position 1 was independent of RBS strength, an important feature for RBS design flexibility and modularity of mARi.

### mARi activity is tuneable via genetic design parameters

To evaluate the features of mARi, we characterised its responsiveness to genetic design parameters and its robustness to different contexts and growth conditions. Two key determinants modulating the repression activity of this post-transcriptional regulator are (i) the degradation rate of RNA complexes and (ii) the ratio of transcript abundance (54). We focused on investigating the relative abundance of mARi in relation to its target mRNA. Unlike previous work where inducible expression was used to modulate relative transcript abundance of sRNA to mRNA target (13, 32, 54), here we constitutively expressed both transcripts to test the impact of different transcript levels, which provides more consistent relative expression. Relative transcript abundance of mARi and mRNA was estimated from the relative promoter strength driving both regulator and target (54). A set of standardised constitutive promoters (P_J23xxx_BASIC_) were used to express a *sfGFP* reporter gene using UTR A-RBSc in a p15A backbone (**Supplementary Fig. 3a**). The functional characterisation results were then used as a basis to infer the relative transcript expression ratio (**Fig. 3a**), which is calculated by dividing the strength of the promoter used for mARi by the promoter expressing the mRNA target. To assess the impact of the mARi:mRNA ratio, a matrix of designs with four different promoter strengths for both mARi and mRNA were constructed (**Fig. 3a, b**). The resulting sfGFP expression showed a consistent trend, with sfGFP fluorescence decreasing as the mARi:mRNA ratio increased (**Fig. 3c, d, Supplementary Fig. 3c, Supplementary Table 3**). The maximum repression was observed when an excess of mARi was present in relation to its target mRNA, presumably because an excess of mARi is required to saturate binding of target mRNA. On the other hand, a low expression ratio led to weak repression, presumably due to an insufficient amount of mARi being available to inactivate mRNA target (30, 54).

**Figure 3.**
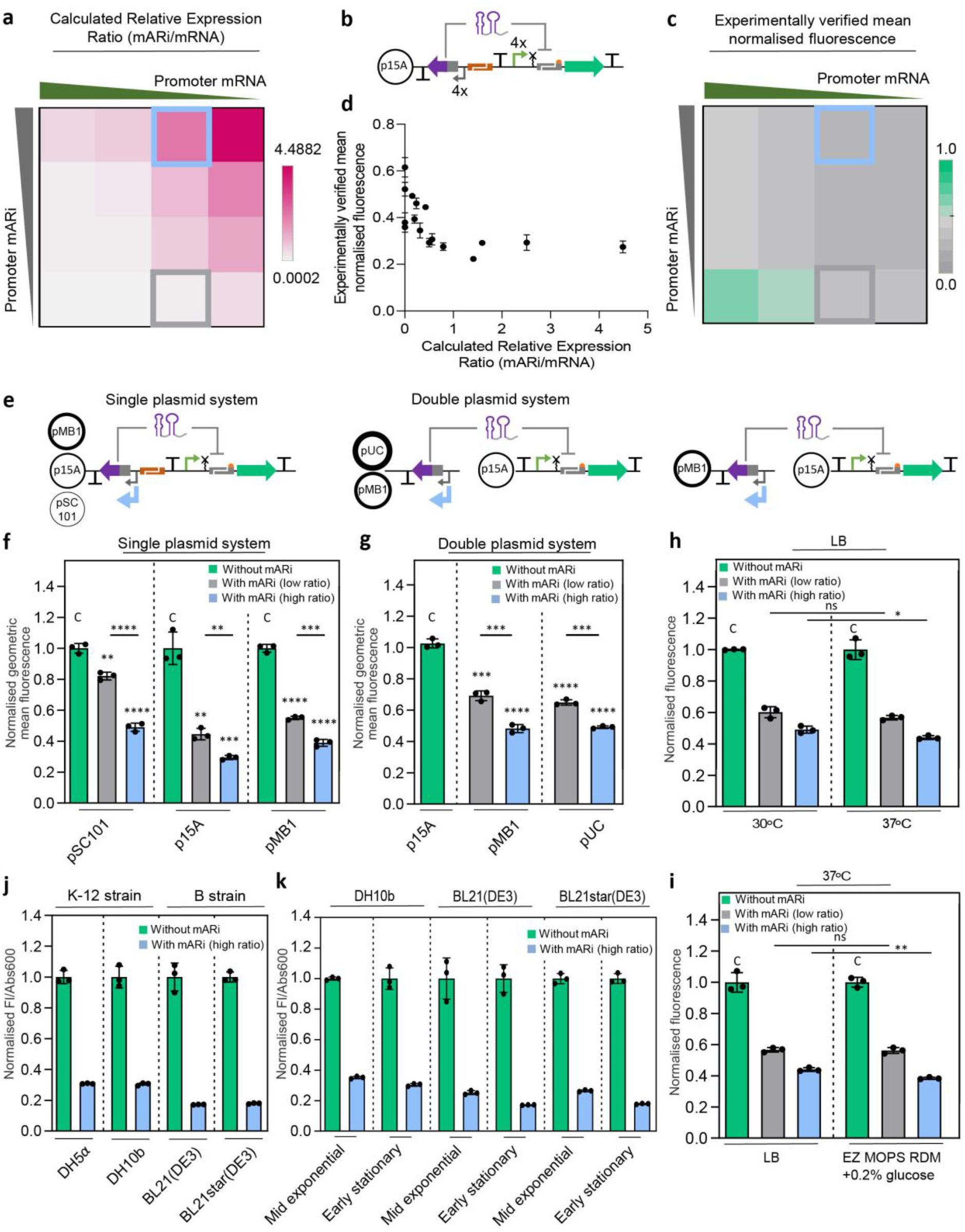
Characterisation of mARi-A across different genetic architectures and growth conditions. **(a)** Heat-map showing relative expression ratio calculated from the relative strength of promoters driving mARi and mRNA expression across a matrix of constructs (**Supplementary Fig. 3b**). The representative conditions used for set-point high and low expression ratios are highlighted with a blue square (P_J23119_-mARi:P_J23101BASIC_-mRNA) and grey square (P_J23101_-mARi:P_J23101BASIC_-mRNA), respectively. **(b)** Schematic of mARi repression constructs used to investigate the effect of transcript ratios using four different strength constitutive promoters driving both mARi and target mRNA in a single plasmid system. **(c**) Heat-map showing the measured relative expression levels (mean normalized fluorescence) of the mARi expression ratio matrix from (**a**). The calculated relative expression ratio and repression activity of mARi are provided in **Supplementary Table 3**. (**d**) Correlation of transcript expression ratios in **a** and normalised fluorescence in **c**. (**e**) Schematic of the mARi-based repression system with high and low expression ratios constructed in single- and double-plasmid systems with different copy numbers. Reporter expression was measured for the single (**f**) and double (**g**) plasmid systems for high and low ratio mARi against a control construct without mARi; sfGFP fluorescence measurements were performed by flow cytometry assay. (**h**) The robustness of the double-plasmid mARi-based regulation in different incubation temperatures. A schematic of the mARi-based repression system in double-plasmid systems used is illustrated. (**i**) The robustness of the double-plasmid mARi-based repression system in a variety of growth media. (**j**) Reporter expression of the single-plasmid high-ratio mARi on p15A was measured in different *E. coli* strains against a control construct without mARi. (**k**) The repression activity of single-plasmid high-ratio mARi on p15A in representative strains (DH10b, BL21 (DE3), and BL21star (DE3)) across different growth phases (**Supplementary Fig. 5**). Assays in **h, i, j** and **k** were performed by plate reader assay. All error bars show the mean ± SD of triplicate measurements (black dots). Statistically significant differences were determined against control without mARi expression (C) using two-tailed Student’s t-test (**** represents p<0.0001, *** represents p<0.001, ** represents p<0.01, * represents p<0.1, ns represents not significant).

**Figure 4.**
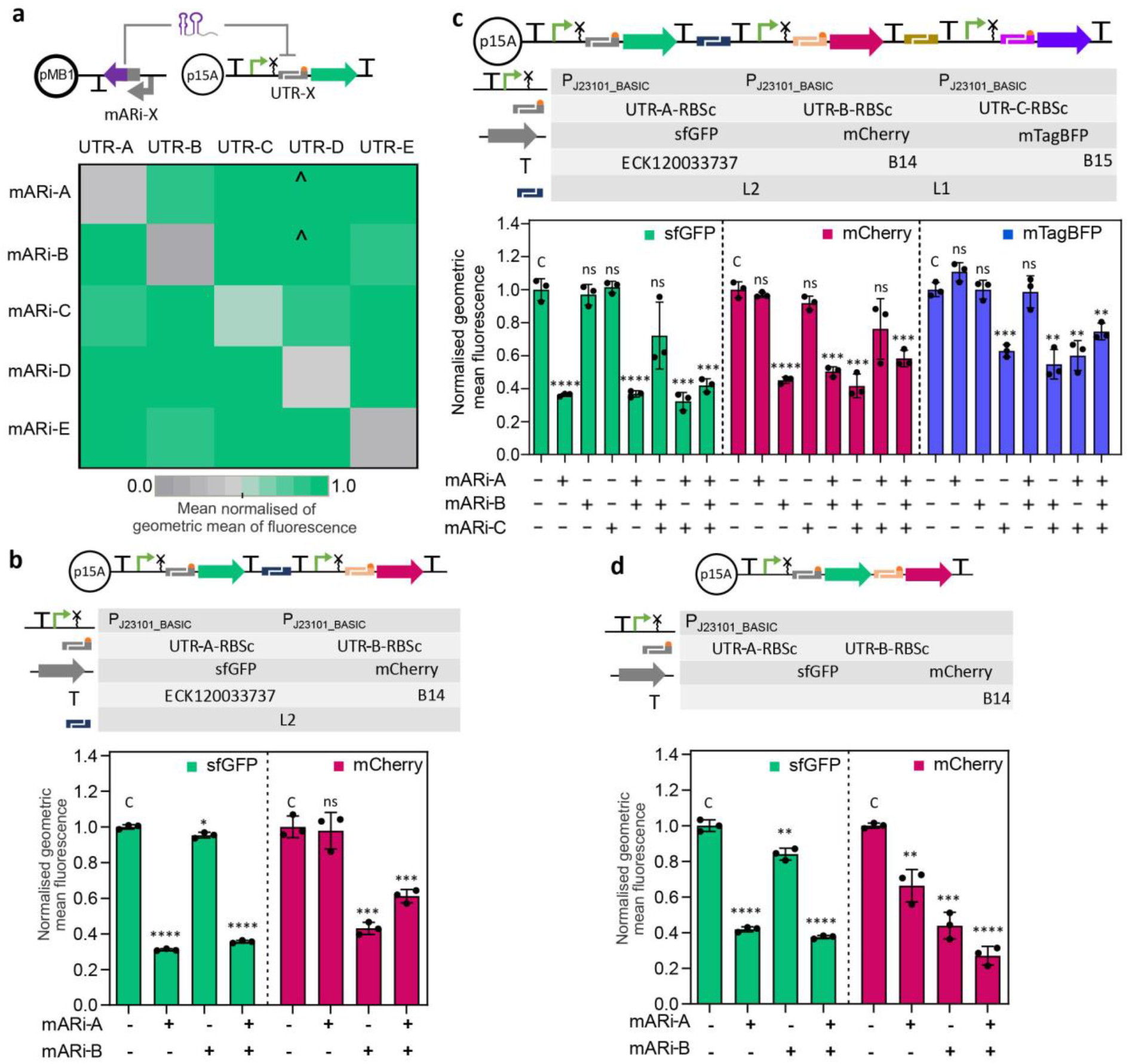
mARi enables multiplex and simultaneous control of gene expression in a multi-gene system. (**a**) A schematic of experimental design to evaluate the target specificity of modular mARi-mRNA pairs is shown. Fluorescence response matrix of all possible mARi/UTR pairs measured by flow cytometry assay with triplicate repeats (**Supplementary Fig. 7d**); ^ represents data where the use of mARi-A and mARi-B increased UTR-D-sfGFP expression relative to control. (**b**) A schematic of reporters in dual transcription units is shown: the fluorescence response of reporters in the double transcriptional unit system when combined with all possible mARi-A and mARi-B regulators was measured by flow cytometry assay. (**c**) A schematic of reporters in triple transcription units is shown: the fluorescence response of reporters in the triple transcriptional unit system when combined with all possible mARi-A, mARi-B and mARi-C regulators was measured by flow cytometry assay. (**d**) A schematic of dual reporters in an operon system is shown: the fluorescence response of dual reporters in the operon system when combined with all possible mARi-A and mARi-B regulators was measured by flow cytometry assay. All error bars show the mean ± SD of triplicate measurements (black dots). Statistically significant differences were determined against control without mARi expression (C) using two-tailed Student’s t-test (**** represents p<0.0001, *** represents p<0.001, ** represents p<0.01, represents p<0.1, ns represents not significant).

A further variable affecting intracellular transcript abundance is plasmid copy number used in the system. Designs with both high (blue box in **Fig. 3a**) and low (grey box in **Fig. 3a**) expression ratios were selected to exemplify different levels of transcript abundance. These systems were first assembled with both mARi and its target in a single plasmid, with varying plasmid copy numbers from low to high: pSC101 (∼5 copies per cell), p15A (∼10 copies per cell) and pMB1 (∼15-20 copies per cell) (55) (**Fig. 3e left panel, Supplementary Fig. 4a**). Expression of mARi in either high or low expression ratio repressed sfGFP expression for all tested plasmid copy numbers, demonstrating that the mARi-based regulatory system is functional at different plasmid copy numbers (**Fig. 3f**).

We next reasoned that by placing the mARi on a second, higher copy vector, repression should be more efficient. To test this hypothesis, the mRNA expression cassette was placed on a p15A backbone, while mARi was placed on the higher copy pMB1 and pUC backbones (∼500-700 copies per cell), with both high- and low-expression ratios (**Fig. 3e right panel, Supplementary Fig. 4b**). In this experimental design, the transcript expression ratio was governed by the combination of plasmid copy number and transcription rate of mARi relative to mRNA. Despite mARi being driven by a stronger promoter and being on a higher copy vector, in all cases the double-plasmid systems all showed less repression than that observed with the corresponding single-plasmid systems (**Fig. 3g**). This suggests a role for spatial organisation of transcripts in post-transcriptional repression, as anticipated from recent theoretical analysis (56). The fact that the sRNA-based regulator has a shorter half-life and stability than mRNA may further reduce the mARi effectiveness when it is used in a double plasmid system with a broader physical transcript distribution.

To investigate the robustness of mARi repression system, we further tested the double plasmid system with both high and low expression ratios in different temperatures and growth media. The repression activity of mARi in low expression ratios was not significantly affected by a combination of different growth temperatures and growth media (**Fig. 3h, 3i**). However, a high expression ratio resulted in a slight increase of mARi repression activity in the use of higher incubation temperature and a defined growth medium.

### mARi-based regulation is robust to different genetic contexts

Unlike other biomolecular regulators (e.g. transcription factors, TALEs, zinc fingers, or CRISPRi) (5, 57, 58), mARi only requires transcriptional activity and endogenous Hfq, which is autoregulated by Hfq binding its own mRNA (59). Thus, the expression of a short sequence for mARi would be expected to have a low cellular burden and not significantly impact host cell growth. To test this, the high transcript ratio for mARi in a p15A backbone and control plasmid without mARi expression were used to compare the effect of mARi overexpression on host fitness. We sought to test this repression system in four commonly used *E. coli* strains: DH5α, DH10b, BL21(DE3), and BL21star(DE3). The high expression of mARi did not affect bacterial growth across different host strains, indicating the mARi-based repression system has a low production cost, and does not impact significantly on host cells (**Supplementary Fig. 5**).

We then sought to evaluate the silencing activity of mARi in these *E. coli* K12 and B strains. The mARi-based gene regulation utilises the native *E. coli* Hfq chaperone and degradosome complex (e.g. RNase E) as helper proteins to regulate gene repression. The intracellular abundance of these components is influenced by several factors, including the genotype and growth phase. For instance, the abundance of Hfq is constant during the exponential growth, but reduces when *E. coli* cells reach stationary phase (47, 60–63).

Functional characterisation of mARi at early stationary phase, 8 h (**Supplementary Fig. 5**), showed about 60-80% repression activity in both *E. coli* K-12 and B strains (**Fig. 3j**), while the silencing activity in the B strains was higher than in the K-12 strains for comparable plasmid copy numbers. (**Supplementary Fig. 6**). The mARi system was active across different phases of cell growth (**Fig. 3k**): the *E. coli* K-12 strain (DH10b cells) showed relatively stable repression activity in both mid-exponential and early stationary growth, whereas a slightly increased repression activity was observed in early stationary phase for the BL21(DE3) and BL21star(DE3) strains.

### Expanding a set of orthogonal mARi regulators through seed sequence modification

We sought to create a set of mARi-based regulators based on its underlying modular design principles coupled with the modular UTR-RBS linkers used in BASIC to construct both expression cassettes and operons. Five different UTR-RBS linkers were identified using R2oDNA Designer (35), whilst preserving the same medium strength RBS (RBSc). Computational evaluation of the five selected UTR (UTR-A-E) sequences demonstrated minimal similarity/identity scores (**Supplementary Fig. 7a)** and high Hamming distance scores (**Supplementary Fig. 7b)** between the set. Low sequence similarity and high Hamming distance of non-cognate pairs is desirable to achieve minimal cross-talk interaction and off-target effects (30). The target 35 bp upstream of the RBS sequence were evaluated for their GC content and free binding energy of the target/seed sequences (**Supplementary Table 4)**. The GC content of the seed sequence is in a range of 31% to 43% and resulted in a free binding energy of - 53.9 kcal/mol to -63.9 kcal/mol, which were in the favourable range for sRNA repression (30, 31). The seed sequences were also predicted to have minimal off-target interactions with three different *E. coli* genomes: *E. coli* MG1655, DH10b, and BL21(DE3) **(Supplementary Table 4)**.

To experimentally validate target specificity, we constructed all possible combinations of mARi and target UTR upstream of *sfgfp*; in each case maintaining the same promoter (**Fig. 4a**). The expression of sfGFP was significantly repressed in the presence of the cognate mARi systems relative to the non-cognate pairings (**Fig. 4a, Supplementary Fig. 7d**), confirming their specificity with reduced cross-reactivity between each pair. However, there is an increased expression in UTR-D-sfGFP when mARi-A and mARi-B were used. The lower Hamming score of UTR D towards UTR A (**Supplementary Fig. 7b**) may contribute to this, but it does not completely explain the effect with mARi-B, or other observations; similar effects were not observed for UTR-E, which similarly has lower Hamming scores for UTR-A, -B and -D (**Supplementary Fig. 7b**).

### Modularity and orthogonality of mARi enables simultaneous and multiplexed gene regulation

After demonstrating that mARi regulators exhibit specific interaction towards cognate UTRs in the context of a single gene, we evaluated their ability to operate in a multiplexed reporter gene design. Initially, this was evaluated with both reporter genes (i.e. *sfgfp* and *mcherry*) organised as separate transcriptional units (**Fig. 4b**). Specific and independent repression by each mARi towards its cognate target was observed, with UTR-A-*sfgfp* being selectively targeted by mARi-A and UTR-B-*mcherry* being targeted by mARi-B (**Fig. 4b** and **Supplementary Fig. 8a**). Further, specific and independent repression was also confirmed when three mARi regulators (mARi-A, mARi-B, and mARi-C) were employed to target UTR-A, UTR-B, and UTR-C preceding the *sfgfp, mcherry*, and *mtagbfp* genes, respectively (**Fig. 4c** and **Supplementary Fig. 8b**).

Two sets of mARi regulators were used to modulate gene expression in a two-gene operon with UTR-A-RBSc driving the expression of *sfgfp*, upstream of UTR-B-RBSc driving the expression of *mcherry*, constitutively expressed by the promoter P_J23101_BASIC_ (**Fig. 4d**). With both genes on a single transcript, the targeted repression for the first gene (*sfgfp*) is strongly affected by its cognate mARi-A, with only a minor affect from mARi-B, which targets the downstream *mcherry*; the downstream *mcherry* is most strongly attenuated by its cognate mARi-B, but it is also significantly attenuated by repression of the upstream *sfgfp* with full attenuation only reached with both mARi-A and B (**Fig. 4d** and **Supplementary Fig. 8c**).

## DISCUSSION

Here we report the development, characterisation, and implementation of a modular post-transcriptional regulation system based on *trans*-encoded sRNAs. The modular design of mARi is derived from the highly composable structure of the natural MicC sRNAs scaffold (17, 27). A potential limitation of our system is that our mARi regulators utilise shared endogenous Hfq chaperone for their repression activity. However, endogenous Hfq is autoregulated by Hfq binding its own mRNA, even when expressed from plasmid-borne genes (59) and the intracellular concentration of available Hfq can adapt rapidly to cellular changes in the RNA pool by facilitated recycling of the protein (64). In addition, deviation from the optimum Hfq concentration can lead to suppression of sRNA activity in *E. coli* (29). Efforts to modulate sRNA activity by exogenous expression of Hfq would therefore require careful balancing of the whole system.

Modification of the seed sequence, whilst retaining the sRNA scaffold, has enabled the reprogramming of natural sRNAs with different targets of interests that do not exist in nature (17, 18, 28). Here, we utilised this modularity for targeted gene repression by directing mARi regulators to standardised UTRs in the translation initiation region of the mRNA target. We demonstrated that targeted repression by binding to standardised sequences in the 5’-UTR upstream of the RBS were independent from both RBS and GOI contexts. The computationally designed orthogonal UTR sequences of the BASIC UTR-RBS linkers proved to be ideal target sites for mARi regulators (**Fig. 2c, e** and **4**). The standardised linker sequences that define the UTR have been computationally generated and validated to ensure their orthogonality in the DNA assembly process, as well as to the *E. coli* host (35). Therefore, by targeting the BASIC DNA linkers, it was anticipated that the mARi would exert orthogonal, post-transcriptional gene regulation with minimal cross-interaction to non-cognate targets, bioparts, plasmid backbones and host genotypes (35). The sequence composition of seed/target sites can be refined to optimise their binding affinity. This approach may result in higher repression and orthogonality. Therefore, it could also be coupled with an improved algorithm to create an sRNA optimised orthogonal sequence for the assembly process using BASIC DNA assembly.

The mARi-based repression of a polycistronic mRNA provides some insight into its mechanism of action. Facilitated degradation of the mRNA would be expected to induce a similar repression level on both genes, even when only one is targeted, but single-gene targeting in an operon led to a differential effect between the operon’s genes. A non-targeted gene downstream of the mARi gene target exhibited significant repression, while a non-targeted gene upstream of the mARi gene target exhibited only slight repression (**Fig. 4d**). This behaviour is consistent with steric hindrance of the ribosome for the UTR-RBS preventing translation initiation being the main mode of action of the mARi sRNA, rather than active degradation. The silencing effects on the downstream *mcherry* gene were also additive for both the gene’s cognate mARi and the upstream gene’s mARi, suggesting that repression at each UTR-RBS position was independent. It further suggests that read-through was possible from the upstream gene, even when the downstream RBS was repressed, but this was occluded when both UTRs were bound by their respective mARi regulators.

The mechanism of action being based on the inhibition of translation initiation rather than active degradation may also be due to the designed mARi regulators and UTR-RBS linkers omitting an RNase E binding site, which is essential for RNase E-dependent cleavage. The elimination of RNase E in the mARi design was anticipated to reduce background degradation of unpaired mARi regulators and improve their cellular half-lives. The exclusion of an RNase E cleavage site in the mARi design may be beneficial for the implementation of an orthogonal repression system in various *E. coli* strains, with a response that is independent of the host genotype. Indeed, the system was observed to function across different *E. coli* strains and growth phases, demonstrating both its portability and robustness, notably with minimal difference between BL21(DE3) and BL21star(DE3), the latter being deficient in RNase E (**Fig. 3**).

The modular nature of the design framework meant facile variation of genetic design parameters. Our observations of design characteristics were consistent with previous studies (30), where repression increased as a function of length and hence free binding energy (**Supplementary Fig. 2**). Importantly, we have also identified spatial organisation and relative transcript abundance as being critical factors in efficient repression by sRNA (**Fig. 3**). Expressing mARi in a high expression ratio and a single plasmid system with medium plasmid copy number (p15A) resulted in up to 80% repression activity. With *trans* expression of mARi from a second plasmid, the repression ratio was always worse, even when it was encoded on a higher plasmid copy number. Also somewhat surprising was the low efficiency of repression in a low-copy (pSC101) single plasmid system. This likely represents the importance of having the correct balance of all components relative to the native Hfq availability, as anticipated from Sagawa et al. (29). The importance of spatial organisation is somewhat surprising given the abundance of natural *trans*-encoded systems, but it is likely to originate from enhanced proximity and hence binding between the mARi and its target.

The modular nature of the mARi design enabled control of different genes and facilitated multiplexing of target genes. In total, five pairs of orthogonal mARi /UTR regulators were created and tested in this work. These mARi regulators were applied to multiplexed and simultaneous post-transcriptional regulation in multi-gene systems, including both multiple transcriptional units and an operon architecture (**Fig. 4**). Importantly, the orthogonal basis of the BASIC design framework removed the need to design or insert bespoke target sites upstream of target genes, a drawback of previous works focused on reusable *trans*-encoded small RNAs (7, 24–26, 28); it was implemented by simply assembling the mARi regulators and their cognate targets using BASIC DNA assembly without re-optimisation of the regulator for different target genes. This extensibility is essential for the simultaneous regulation of multiple genes in metabolic engineering, layered genetic control, and advanced genetic circuit applications, while scalability is possible through our recently demonstrated low cost automation platform for BASIC assembly (33).

## Author Contributions

A.D.: Designed and performed all experiments and data analysis, devised the experimental strategy and wrote the manuscript.

M.S.: Devised experimental strategy and design, assisted with the experiments and data interpretation and wrote the manuscript.

G.-B.S.: Contributed to the experimental strategy, assisted with experimental design and data interpretation and wrote the manuscript.

G.S.B.: Contributed to the experimental strategy, assisted with the experimental design and data interpretation and wrote the manuscript.

## Acknowledgements

A.D. received a PhD scholarship from Indonesia Endowment Fund for Education (LPDP). G.-B.S. gratefully acknowledges the support of the UK EPSRC through the EPSRC Fellowship for Growth EP/M002187/1 and of the Royal Academy of Engineering through the Chair in Emerging Technology programme. G.S.B. acknowledges the support of UK Research and Innovation through the Engineering and Physical Sciences Research Council [EP/R034915/1]; as well as the EU through H2020 [820699].

