## Supplementary material for "A modular RNA interference system for multiplexed gene regulation": Supp Dwijayanti et al_NAR_Final_Rev2.pdf

Supplementary Fig. 1. Predicted base-pairing interactions of the designed mARi and mRNA target.
Supplementary Fig. 2. Functional characterisation of mARi-based gene regulation targeting position 1 with different lengths
Supplementary Fig. 3. Characterisation of constitutive promoters and transcript stoichiometry.
Supplementary Fig. 4. Measured sfGFP reporter of mARi-based gene regulation for the single and double plasmid systems.
Supplementary Fig. 5. Growth profile and fluorescence output of different host strains expressing the mARi-based repression system.
Supplementary Fig. 6. mARi-based regulation in different *E. coli* strains.
Supplementary Fig. 7. Orthogonal repression by mARi.
Supplementary Fig. 8. Multiplexed and simultaneous gene regulation by mARi.
Supplementary Fig. 9. Maps for the plasmids used in this study.

Supplementary Table 1. Summary of mARi variants used to evaluate the impact of the position of the target site.
Supplementary Table 2. UTR-RBS BASIC linker sequences used for testing the performance of mARi-A when combined with different RBS.
Supplementary Table 3. The calculated relative expression ratio and repression activity of mARi.
Supplementary Table 4. Orthogonal mARis and off-target prediction towards *E. coli* genome.
Supplementary Table 5. List of standardised bioparts sequences used in this study.
Supplementary Table 6. List of orthogonal BASIC linker sequences used in this study.

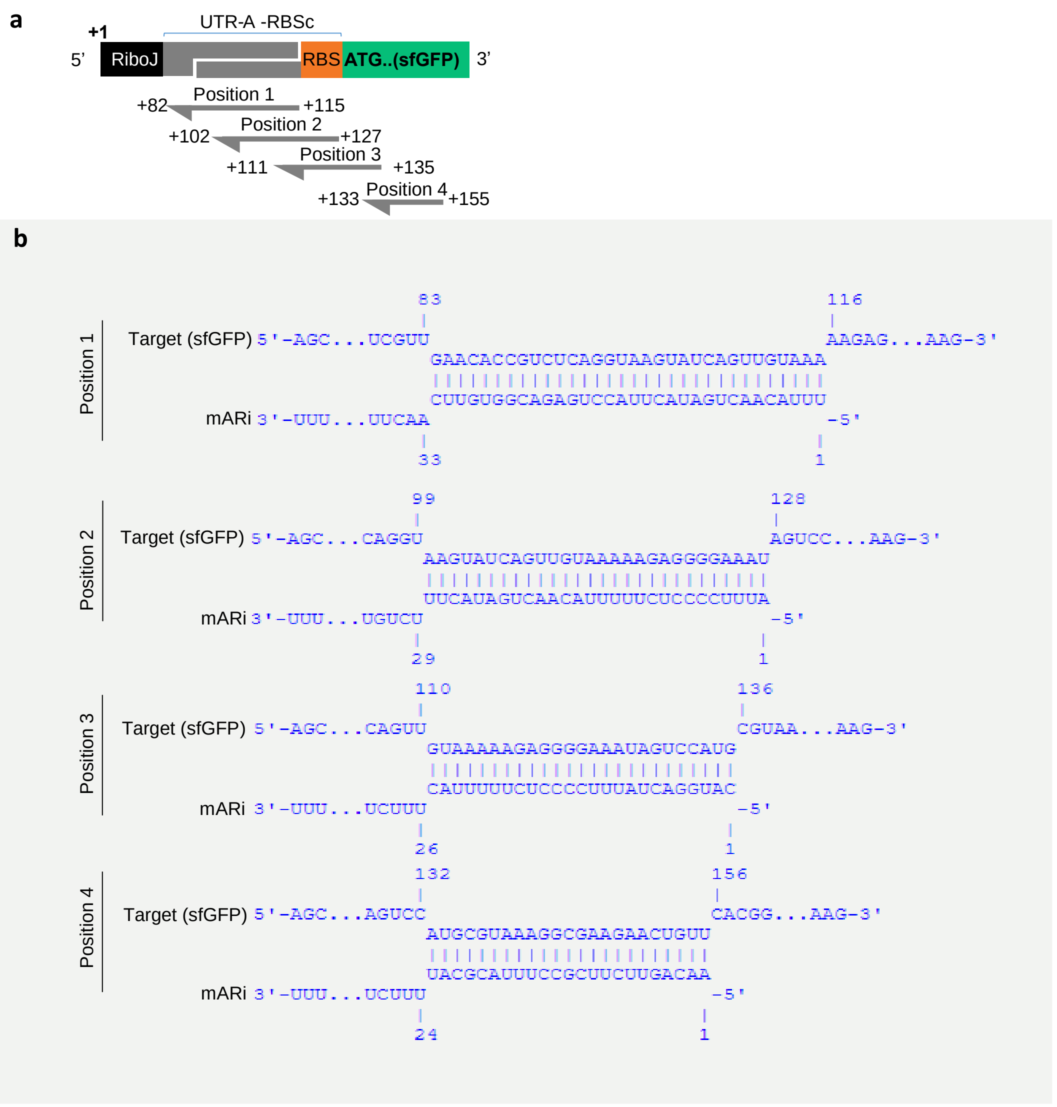


**Supplementary Fig. 1. Predicted base-pairing interactions of the designed mARi and mRNA target.**

(**a**) Schematic design of target site selection (Positions 1-4) of mARi-mediated repression. Arrows show the direction of the reverse complementary sequence in the mARi design. (**b**) Predicted base pairing of mARi-A with mRNA targets (sfGFP) for the four different target positions. Numbers indicate the relative positions of bases from the Transcription Start Site (+1).

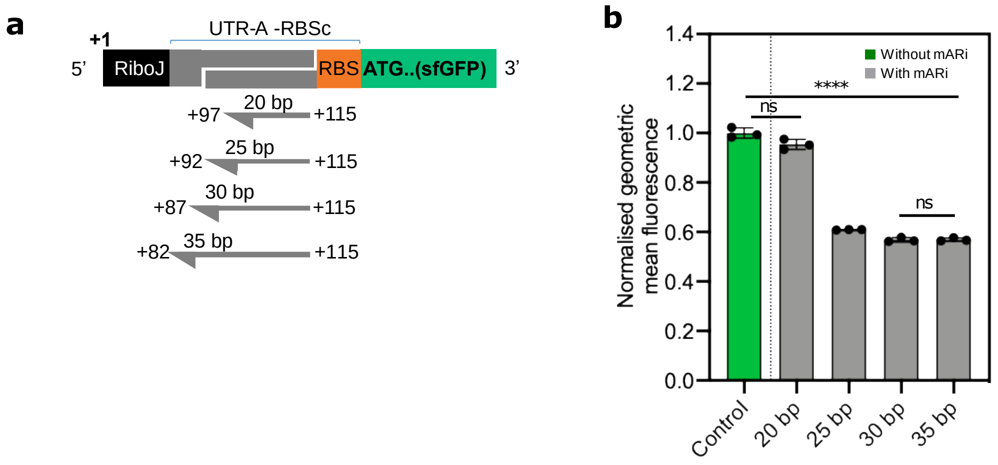


**Supplementary Fig. 2. Functional characterisation of mARi-based gene regulation targeting position 1 with different lengths.**

(**a**) Schematic design of mARi-mediated repression targeting position 1 with different lengths: 20, 25, 30, and 35 bp (Supplementary Table 1). Arrows show the direction of the reverse complementary sequence in the mARi design. (**b**) Normalised fluorescence of mARi-based gene regulation targeting position 1 with different lengths. Data are shown with error bars for the mean ± SD of triplicate measurements (black dots). Statistically significant differences were determined using two-tailed Student's t-test (**** represents $p<0.0001$ and ns represents not significant).

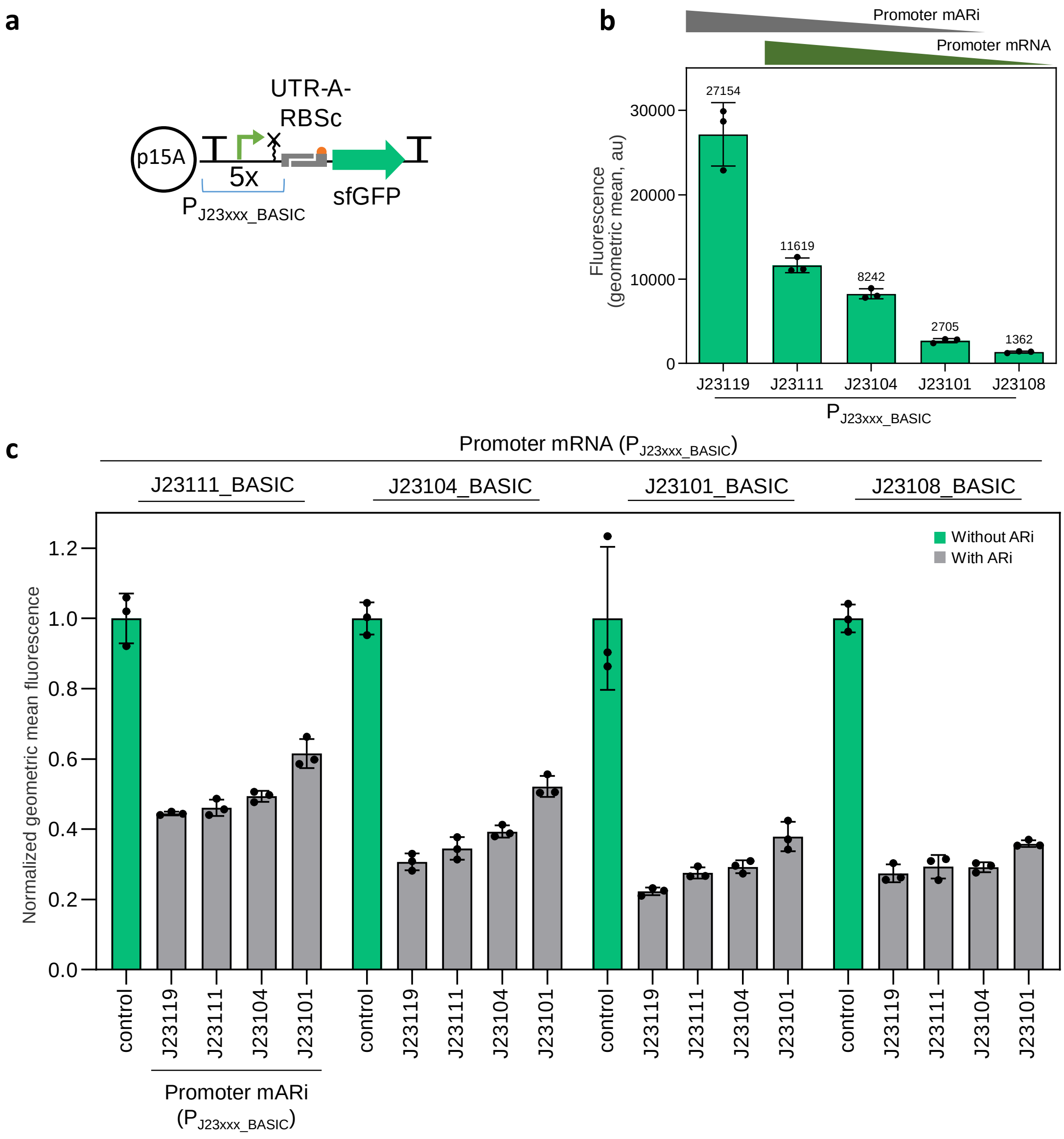


**Supplementary Fig. 3. Characterisation of constitutive promoters and transcript stoichiometry.**

(**a**) Schematic of the expression system used to characterise our standardised constitutive promoter set ($P_{J23xxx\_BASIC}$) to express a *sfGFP* reporter gene using UTR A-RBSc in a p15A backbone. (**b**) Promoter activity for the set of the standardised constitutive promoters. This promoter activity was used to calculate the relative expression ratio (mARi:mRNA) in **Fig. 3a**. The subsets of promoters used for mRNA and mARi are indicated by the coloured wedges. (**c**) Normalised fluorescence from flow cytometry assay for all constructs used to evaluate transcript stoichiometry in **Fig. 3c**. Data are shown as the mean ± SD of three biological repeats (black dots). The calculated relative expression ratio and repression activity of mARi are provided in **Supplementary Table 3**.

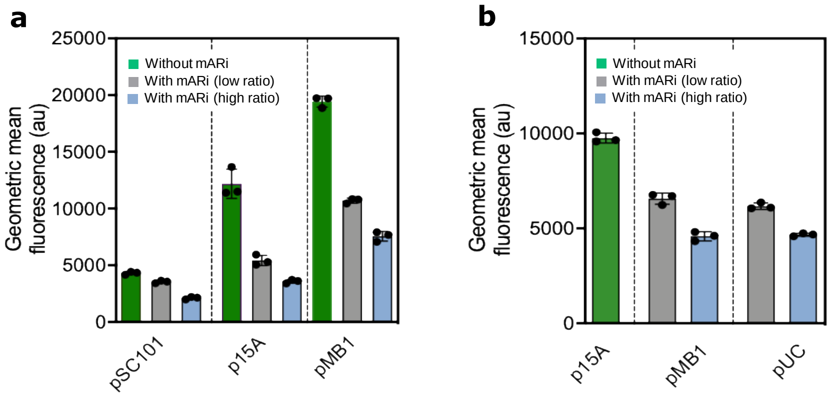


**Supplementary Fig. 4. Measured sfGFP reporter of mARi-based gene regulation for the single and double plasmid systems.**

Measured reporter expression for the single (**a**) and double (**b**) plasmid systems for high and low ratio mARi against a control construct without mARi. The sfGFP fluorescence measurements were performed by flow cytometry assay.

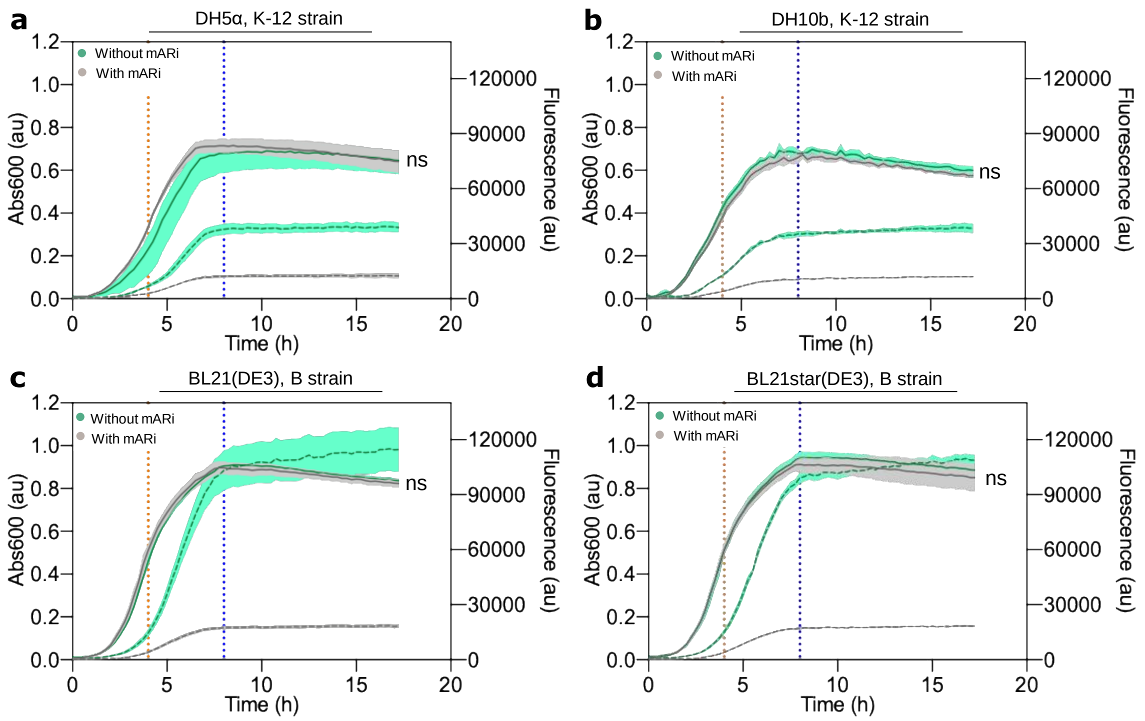


**Supplementary Fig. 5. Growth profile and fluorescence output of different host strains expressing the mARi-based repression system.**

Growth curves (solid lines) and fluorescence outputs (dashed lines) of strains with and without mARi expression in different host strains from a plate reader assay: (**a**) DH5α (K-12 strain), (**b**) DH10b (K-12 strain), (**c**) BL21(DE3) (B strain) and (**d**) BL21star(DE3) (B strain). Data associated with **Fig. 3g** were taken during early stationary phase (blue dotted line); Data associated with **Fig. 3h** were taken during mid-exponential phase at around 4h (orange dotted line) and early stationary phase at around 8h (blue dotted line). Lines show the mean from 3 independent measurements with the shaded area showing ± SD. Statistically significant differences were determined from independent data points at the indicated time points using two-tailed Student's t-test (ns used to denote "not significant").

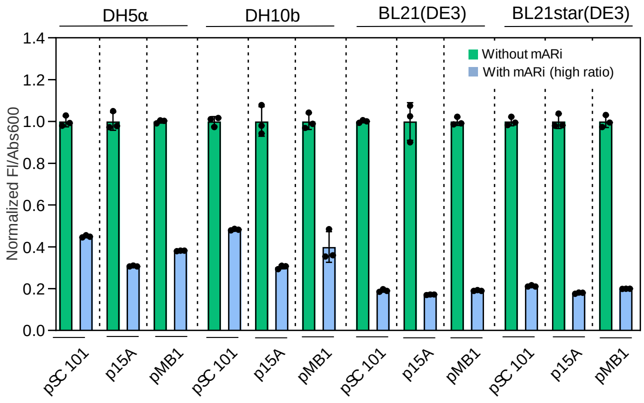


**Supplementary Fig. 6. mARi-based regulation in different *E. coli* strains.**

The performance of mARi repression with a high expression ratio (mARi>mRNA), single plasmid system in four host strains was measured using a plate reader assay. Strains DH5α, DH10b, BL21(DE3), and BL21star(DE3) were used, with three different plasmid copy numbers: pSC101, p15A, and pMB1. Data were taken during early stationary phase (8 h). Data associated with **Fig. 3g, h** were taken from the p15A backbone. Data are shown as the mean ± SD of three independent repeats (black dots).

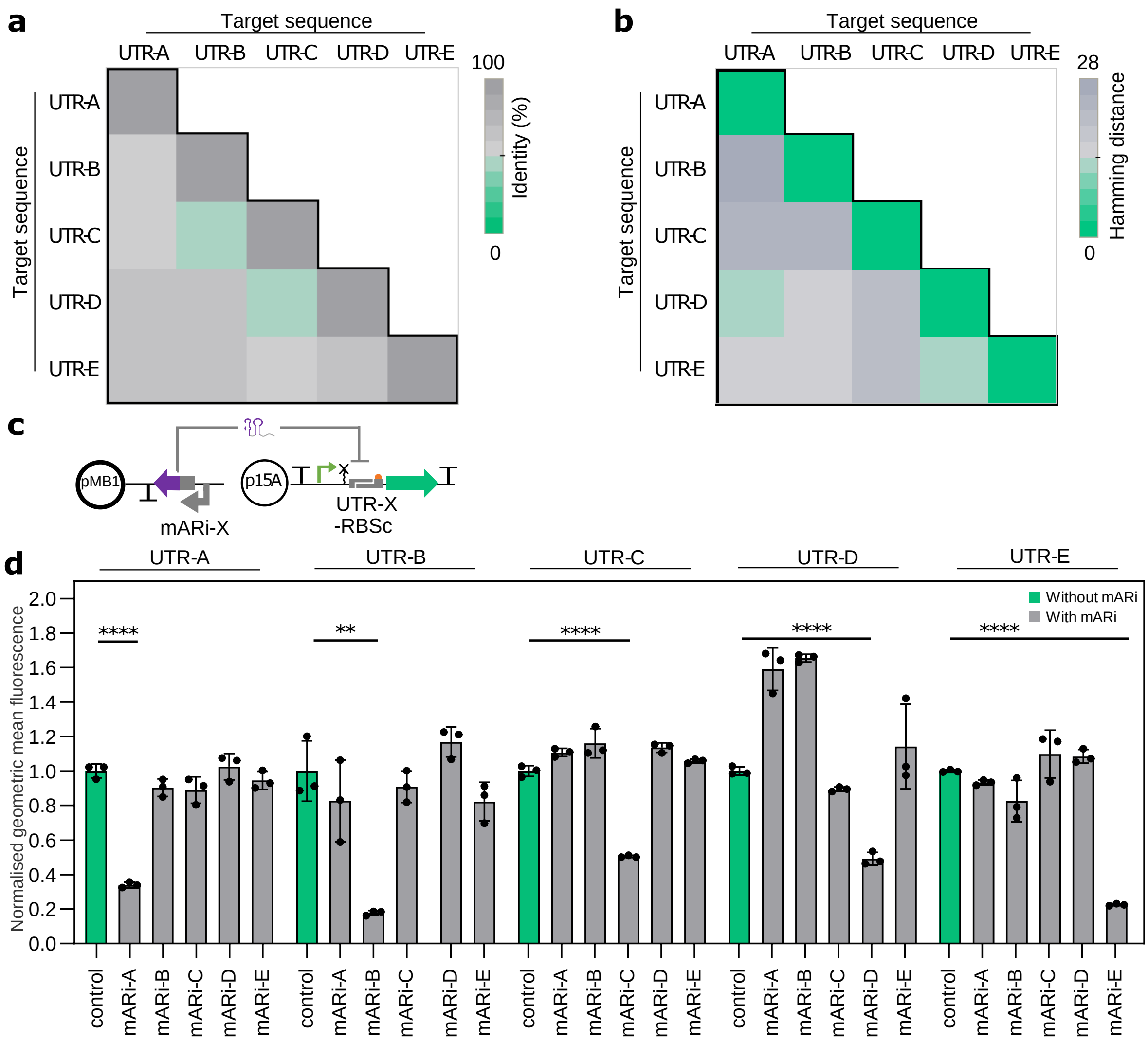


**Supplementary Fig. 7. Orthogonal repression by mARi.**

(**a**) The calculated identity similarities of each target sequence in UTR pairs was obtained using the EMBOSS needle method (https://www.ebi.ac.uk/Tools/psa/emboss_needle/). (**b**) The calculated Hamming distance of each target sequence in UTR pairs. (**c**) Schematic of the genetic constructs used to evaluate target specificity of modular mARi-X and UTR-X pairs. The mRNA expression cassettes driven by $P_{J23101\_BASIC}$ were located in the p15A backbone while the mARi expression cassettes with $P_{J23119}$ were cloned in a pMB1 backbone. Both expression plasmids were then co-transformed into DH5α cells. (**d**) Response of all mARi and target sequence combinations were measured by flow cytometry assay after 6 h of incubation, and the fluorescence was normalised to control cells with sfGFP and without mARi. Data are shown as the mean ± SD of three independent repeats (black dots). Statistically significant differences were determined using two-tailed Student's t-test (**** represents p<0.0001, ** represents p<0.01).

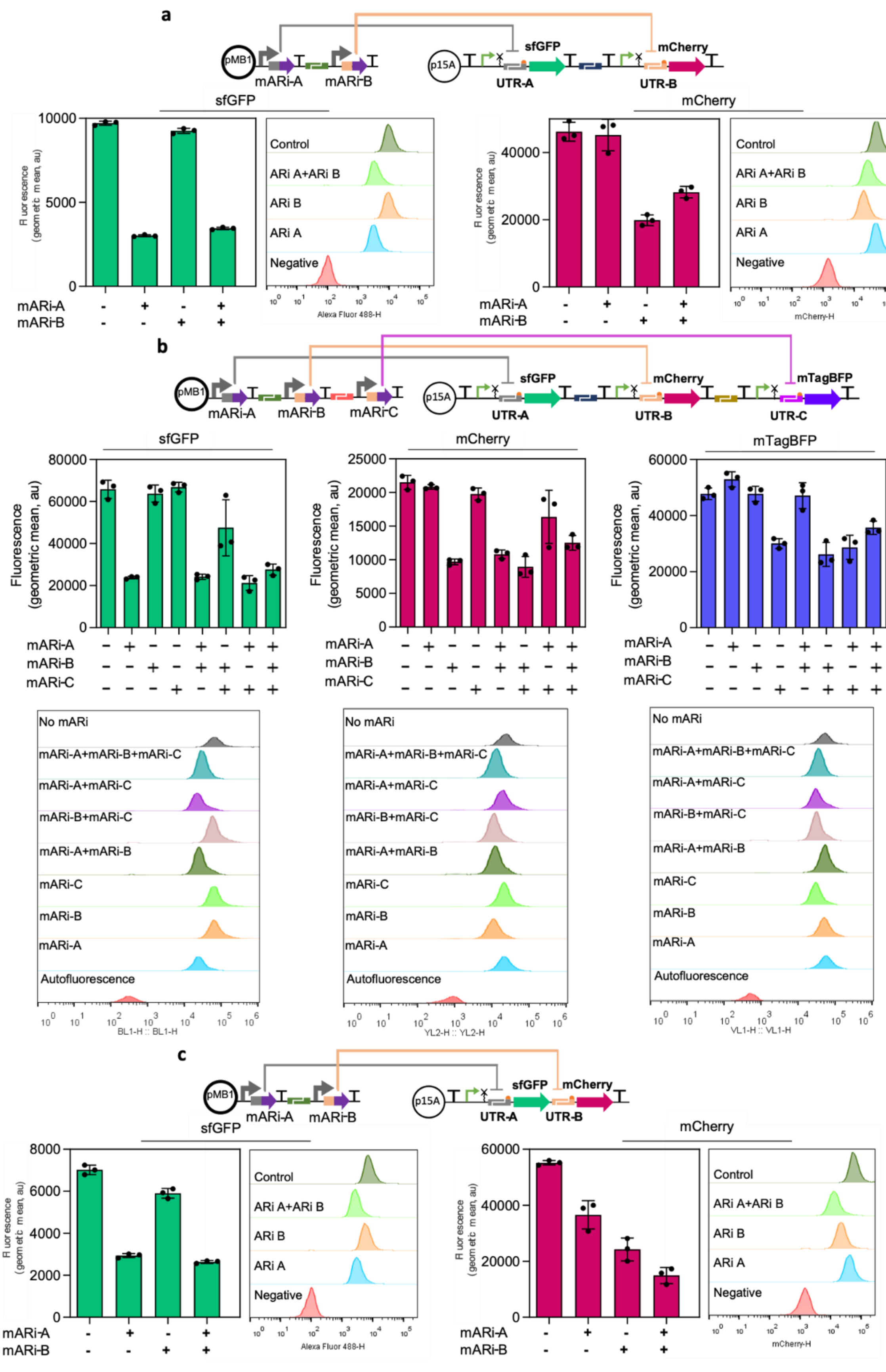


**Supplementary Fig. 8. Multiplexed and simultaneous gene regulation by mARi.**

Measuring the response of mARi regulation in multi-gene systems: independent transcriptional units of two-gene (**a**) and three-gene (**b**), and of operon constructs (**c**). In each case sfGFP was downstream of UTR-A-RBSc, mCherry was downstream of UTR-B-RBS-c, and mTagBFP was downstream UTR-C-RBSc. mARi-A, mARi-B, and mARi-C were expressed both separately and together and their effect on fluorescent protein output was measured by flow cytometry assay. Data are shown as the mean ± SD of three biological repeats (black dots); flow cytometry histograms are also shown.

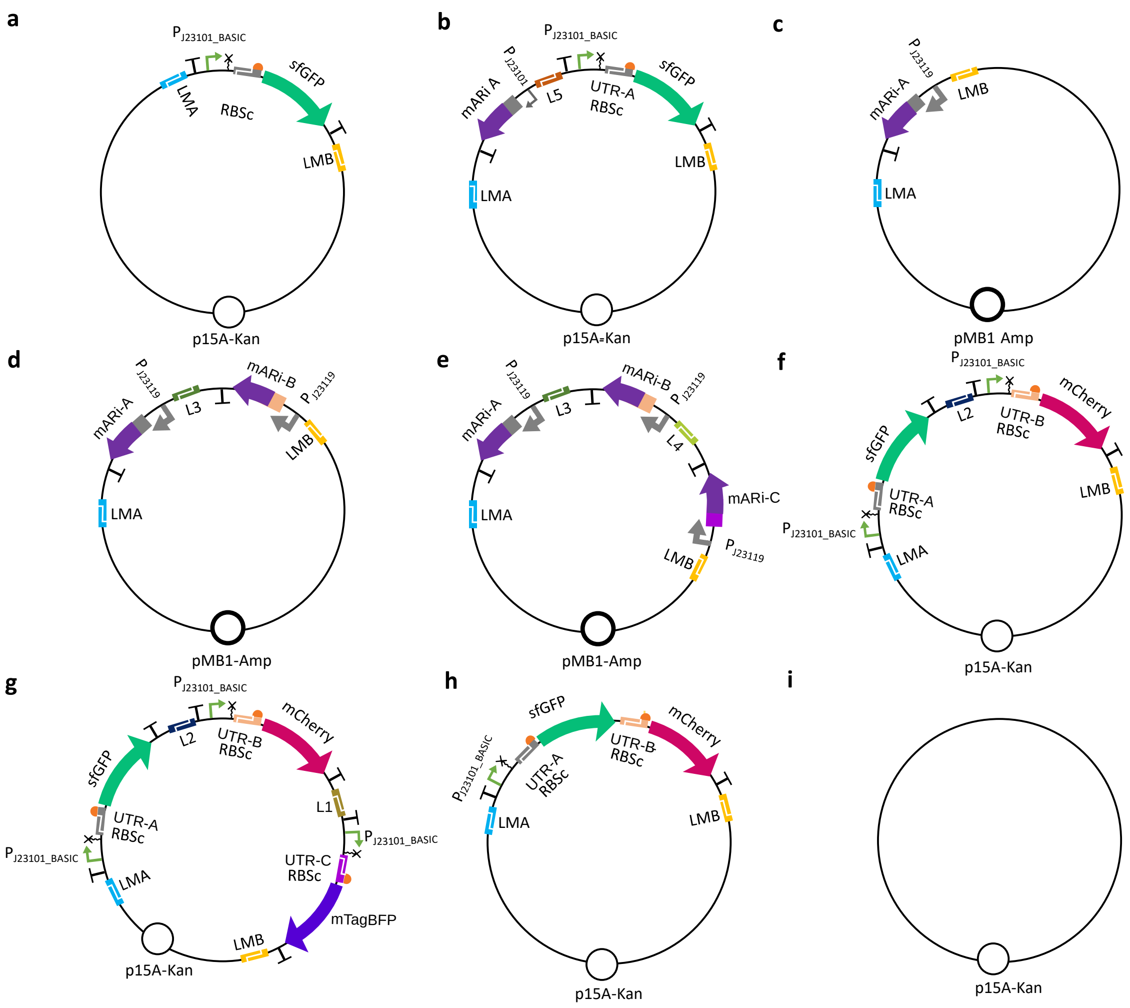


**Supplementary Fig. 9. Maps for the plasmids used in this study.**

(**a**) Map of a plasmid containing a single transcriptional unit of the UTR-A-RBSc-sfGFP expression cassette. (**b**) Map of a plasmid harbouring the $P_{J23101}$ mARi repression system that was used for identifying target site position, transcript stoichiometry, and spatial organisation. (**c**) Map of a single mARi expression plasmid. (**d**) Map of a double mARi expression plasmid. (**e**) Map of a triple mARi expression plasmid. (**f**) Map of a plasmid containing dual transcriptional units for UTR-A-RBSc-sfGFP and UTR-B-RBSc-mCherry. (**g**) Map of a plasmid containing triple transcriptional units for UTR-A-RBSc-sfGFP, UTR-B-RBSc-mCherry, and UTR-C-RBSc-mTagBFP. (**h**) Map of a plasmid containing an operon system with UTR-A-RBSc sfGFP and UTR-B-RBSc mCherry. (**i**) Map of an empty plasmid containing an origin of replication and antibiotic resistance genes. Details of BASIC bioparts used to build the plasmids are provided in **Supplementary Table 5**. The list of orthogonal linkers used to construct the plasmids using BASIC is provided in **Supplementary Table 6**.

**Supplementary Table 1. Summary of mARi variants used to evaluate the impact of the position of the target site.**

| Parameter | mARi-A targeting Position 1 (20 bp) | mARi-A targeting Position 1 (25 bp) | mARi-A targeting Position 1 (30 bp) | mARi-A targeting Position 1 (35 bp) | mARi-A targeting Position 2 | mARi-A targeting Position 3 | mARi-A targeting Position 4 |
|---|---|---|---|---|---|---|---|
| **Context dependency**[a] | | | | | | | |
| Specific to the upstream region of UTR sequence | Yes* | Yes* | Yes* | Yes* | Yes* | Yes* | No |
| Specific to the RBS sequence | No* | No* | No* | No* | Yes | Yes | No* |
| Specific to the GOI sequence | No* | No* | No* | No* | No* | No* | Yes |
| **Target sequence**[b] | 5’-AGGTAAGTATCAGTTGTAAA-3’ | 5’-GTCTCAGGTAAGTATCAGTTGTAAA-3’ | 5’-ACACCGTCTCAGGTAAGTATCAGTTGTAAA-3’ | 5’-GTTGAACACCGTCTCAGGTAAGTATCAGTTGTAAA-3’ | 5’-GTATCAGTTGTAAAAAGAGGGGAAAT-3’ | 5’-GTAAAAAGAGGGGAAATAGTCCATG-3’ | 5’-ATGCGTAAAGGCGAAGAACTGTT-3’ |
| **GC content (%)** | 30 | 36 | 40 | 40 | 34.62 | 40 | 43.8 |
| **Length (bp)** | 20 | 25 | 30 | 35 | 26 | 25 | 23 |
| **Free binding energy (kcal/mol)**[c] | -28.6 | -40.3 | -52 | -61 | -40.7 | -41.4 | -38.6 |
| **mARi sequence**[d] | TTTACAACTGATACTTACCTtttctgttgggccattgcattgccactgattttccaacatataaaaagacaagcccgaacagtcgtccgggcttttttt | TTTACAACTGATACTTACCTGAGACtttctgttgggccattgcattgccactgattttccaacatataaaaagacaagcccgaacagtcgtccgggcttttttt | TTTACAACTGATACTTACCTGAGACGGTGTtttctgttgggccattgcattgccactgattttccaacatataaaaagacaagcccgaacagtcgtccgggcttttttt | TTTACAACTGATACTTACCTGAGACGGTGTTCAACtttctgttgggccattgcattgccactgattttccaacatataaaaagacaagcccgaacagtcgtccgggcttttttt | ATTTCCCCTCTTTTTACAACTGATACtttctgttgggccattgcattgccactgattttccaacatataaaaagacaagcccgaacagtcgtccgggcttttttt | CATGGACTATTTCCCCTCTTTTTACtttctgttgggccattgcattgccactgattttccaacatataaaaagacaagcccgaacagtcgtccgggcttttttt | AACAGTTCTTCGCCTTTACGCATtttctgttgggccattgcattgccactgattttccaacatataaaaagacaagcccgaacagtcgtccgggcttttttt |

[a] The dependency of the target site position towards different contexts is highlighted. * denotes highly desirable for modularity. [b] Target sequence within the UTR-RBS BASIC linker (1) used in a sfGFP expression cassette. [c] The free binding energy of the base-pairing region was calculated at 37˚C (see Methods). [d] Seed sequences are shown in uppercase, while the MicC sRNA scaffold is shown in lowercase.

**Supplementary Table 2. UTR-RBS BASIC linker sequences used for testing the performance of mARi-A when combined with different RBS.**

| UTR-RBS linker | Sequence[a] | Mean relative expression | Predicted strength | |
|---|---|---|---|---|
| | | | **RBS calculator v2.0** (2, 3) | **EMOPEC** (4) |
| UTR-A-RBSa | ggctcgttgaacaccgtctcaggtaagtatc agttgtaa**aaagaggagaaa**tagtcc | 1.00 | 715277.31 | 77.8% |
| UTR-A-RBSb | ggctcgttgaacaccgtctcaggtaagtatc agttgtaa**atctaaggaggt**agtcc | 0.52 | 673992.76 | 80.5% |
| UTR-A-RBSc | ggctcgttgaacaccgtctcaggtaagtatc agttgtaa**aaagaggggaaa**tagtcc | 0.35 | 689020.88 | 79.5% |
| UTR-A-RBSd | ggctcgttgaacaccgtctcaggtaagtatc agttgtaa**atcccaggaggt**agtcc | 0.23 | 17315.58 | 90.5% |
| UTR-A-RBSe | ggctcgttgaacaccgtctcaggtaagtatc agttgtaa**atctcgggaggt**agtcc | 0.18 | 715277.31 | 77.3% |

Predicted output values are shown for computational evaluation of the "strength" of UTR-RBS sequences including the first 100 bases of the sfGFP reporter DNA sequence using two different methods (RBS calculator v2.0 and EMOPEC).

[a] DNA sequence colours correspond to upstream scar (blue) and downstream scar (orange). The DNA sequence in underlined-bold indicates the RBS sequence.

**Supplementary Table 3. The calculated relative expression ratio and repression activity of mARi.**

| mARi promoter | mRNA promoter | Calculated relative expression ratio (mARi/mRNA) | Experimentally verified normalized fluorescence |
|---|---|---|---|
| J23119 | J23111 | 0.424654 | 0.444787 ± 0.004998 |
| J23111 | J23111 | 0.237425 | 0.461128 ± 0.023085 |
| J23104 | J23111 | 0.150723 | 0.493309 ± 0.015242 |
| J23101 | J23111 | 0.000199 | 0.615604 ± 0.041495 |
| J23119 | J23104 | 0.561215 | 0.306795 ± 0.024429 |
| J23111 | J23104 | 0.313776 | 0.344862 ± 0.031806 |
| J23104 | J23104 | 0.199192 | 0.393503 ± 0.017245 |
| J23101 | J23104 | 0.000263 | 0.521379 ± 0.029825 |
| J23119 | J23101 | 1.410095 | 0.222778 ± 0.010729 |
| J23111 | J23101 | 0.788385 | 0.275617 ± 0.016233 |
| J23104 | J23101 | 0.500486 | 0.292916 ± 0.018146 |
| J23101 | J23101 | 0.000662 | 0.378971 ± 0.041516 |
| J23119 | J23108 | 4.48818 | 0.274005 ± 0.025352 |
| J23111 | J23108 | 2.509346 | 0.29306 ± 0.033318 |
| J23104 | J23108 | 1.592994 | 0.291721 ± 0.013977 |
| J23101 | J23108 | 0.002107 | 0.358764 ± 0.00956 |

**Supplementary Table 4. Orthogonal mARis and off-target prediction towards *E. coli* genome.**

| mARi | Seed sequence[a] (35bp) | GC content of seed sequence | Free binding energy of seed sequence[b] (kcal/mol) | mARi sequence[c] | Off-target prediction[d] | | |
|---|---|---|---|---|---|---|---|
| | | | | | Rank | Free binding energy of base-pairing interaction (kcal/mol) | Gene annotation |
| mARi-A | TTTACAACTGATACTTACCTGAGACGGTGTTCAAC | 40 % | -61 | TTTACAACTGATACTTACCTGAGACGGTGTTCAACtttctgttgggccattgcattgccactgattttccaacatataaaaagacaagcccgaacagtcgtccgggcttttttt | 1 | -27.95 | RNA pyrophosphohydrolase |
| | | | | | 2 | -28.08 | YhcH/YigK/YiaL family protein |
| | | | | | 3 | -25.69 | 16S rRNA (cytosine(1407)-C(5))-methyltransferase RsmF |
| mARi-B | TTACAATAGATTTTACCGTCAGACCACGAGATACC | 40 % | -61 | TTACAATAGATTTTACCGTCAGACCACGAGATACCtttctgttgggccattgcattgccactgattttccaacatataaaaagacaagcccgaacagtcgtccgggcttttttt | 1 | -33.85 | DUF386 family protein cupin superfamily |
| | | | | | 2 | -25.10 | 16S rRNA m(5)C1407 methyltransferase SAM-dependent |
| | | | | | 3 | -19.07 | DUF554 family putative inner membrane protein |
| mARi-C | TTTTCTGCTACCCTTATCTCAGCCAATAGTAACAC | 40 % | -62 | TTTTCTGCTACCCTTATCTCAGCCAATAGTAACACtttctgttgggccattgcattgccactgattttccaacatataaaaagacaagcccgaacagtcgtccgggcttttttt | 1 | -27.42 | NADH:ubiquinone oxidoreductase membrane subunit J |
| | | | | | 2 | -27.04 | Sensory histidine kinase in two-component system with NarP |
| | | | | | 3 | -27.03 | DUF386 family protein cupin superfamily |
| mARi-D | TTTACATAGAATACACAGCCGGGACAGGGTATAAC | 42.86 % | -63.9 | TTTACATAGAATACACAGCCGGGACAGGGTATAACtttctgttgggccattgcattgccactgattttccaacatataaaaagacaagcccgaacagtcgtccgggcttttttt | 1 | -27.06 | 16S rRNA m(5)C1407 methyltransferase SAM-dependent |
| | | | | | 2 | -26.28 | DUF386 family protein cupin superfamily |
| | | | | | 3 | -23.62 | O-antigen capsule forming protein-tyrosine-phosphatase;Etk-P dephosphorylase |

| | | | | | | | |
|---|---|---|---|---|---|---|---|
| mARi-E | TTTACATATGTTTTATCGTCAAGACGCTGTATAAC | 31.43 % | -53.9 | TTTACATATGTTTTATCGTCAAGACGCTGTATAACtttctgttgggccattgcattgccactgattttccaacatataaaaagacaagcccgaacagtcgtccgggcttttttt | 1 | -28.14 | DUF386 family protein cupin superfamily |
| | | | | | 2 | -23.97 | Enterobactin/ferrienterobactin esterase |
| | | | | | 3 | -21.20 | 16S rRNA m(5)C1407 methyltransferase SAM-dependent |

[a] Seed sequence in the mARi cassette is the reverse complementary sequence for the cognate target sites within the UTR-RBS BASIC linker (1). [b] The free binding energy of the base-pairing region was calculated at 37˚C (see Methods). [c] Seed sequences are shown in uppercase, while the MicC sRNA scaffold is shown in lowercase. [d] The off-target effect of full sequence of mARis was computed using CopraRNA (5–7) against NC_000913 (*E. coli* MG1655), NC_010473 (*E. coli* DH10b), NC_012892 and NC_012971 (*E. coli* BL21(DE3)) as host references (see Methods).

**Supplementary Table 5. List of standardised bioparts sequences used in this study.**

| Part name | Type | Sequence[a] (5'→3') | Genbank Accession Number |
|---|---|---|---|
| $P_{J23119\_BASIC}$ | Standardised constitutive promoter[b] | tctggtgggtctctgtccCCAATTATTGAACACCCTTCGGGGTGTTTTTTTGTTTCTGGTCTACCATCTCGTTGTGATAAATA GACCTGAAGTGCCTACTCTGGAAAATCTTTGACAGCTAGCTCAGTCCTAGGTATAATGCTAGCAGCTGTCACCGG ATGTGCTTTCCGGTCTGATGAGTCCGTGAGGACGAAACAGCCTCTACAAATAATTTTGTTTAAggctcgggagacctat cg | MT904005 |
| $P_{J23111\_BASIC}$ | Standardised constitutive promoter[b] | tctggtgggtctctgtccCCAATTATTGAACACCCTTCGGGGTGTTTTTTTGTTTCTGGTCTACCATCTCGTTGTGATAAATA GACCTGAAGTGCCTACTCTGGAAAATCTTTGACGGCTAGCTCAGTCCTAGGTATAGTGCTAGCAGCTGTCACCGG ATGTGCTTTCCGGTCTGATGAGTCCGTGAGGACGAAACAGCCTCTACAAATAATTTTGTTTAAggctcgggagacctat cg | MT904006 |
| $P_{J23104\_BASIC}$ | Standardised constitutive promoter[b] | tctggtgggtctctgtccCCAATTATTGAACACCCTTCGGGGTGTTTTTTTGTTTCTGGTCTACCATCTCGTTGTGATAAATA GACCTGAAGTGCCTACTCTGGAAAATCTTTGACAGCTAGCTCAGTCCTAGGTATTGTGCTAGCAGCTGTCACCGG ATGTGCTTTCCGGTCTGATGAGTCCGTGAGGACGAAACAGCCTCTACAAATAATTTTGTTTAAggctcgggagacctat cg | MT904007 |
| $P_{J23101\_BASIC}$ | Standardised constitutive promoter[b] | tctggtgggtctctgtccCCAATTATTGAACACCCTTCGGGGTGTTTTTTTGTTTCTGGTCTACCATCTCGTTGTGATAAATA GACCTGAAGTGCCTACTCTGGAAAATCTTTTACAGCTAGCTCAGTCCTAGGTATTATGCTAGCAGCTGTCACCGG ATGTGCTTTCCGGTCTGATGAGTCCGTGAGGACGAAACAGCCTCTACAAATAATTTTGTTTAAggctcgggagacctat cg | MT904008 |
| $P_{J23108\_BASIC}$ | Standardised constitutive promoter[b] | tctggtgggtctctgtccCCAATTATTGAACACCCTTCGGGGTGTTTTTTTGTTTCTGGTCTACCATCTCGTTGTGATAAATA GACCTGAAGTGCCTACTCTGGAAAATCTCTGACAGCTAGCTCAGTCCTAGGTATAATGCTAGCAGCTGTCACCGG ATGTGCTTTCCGGTCTGATGAGTCCGTGAGGACGAAACAGCCTCTACAAATAATTTTGTTTAAggctcgggagacctat cg | MT904009 |
| $P_{J23101}$_mARi-A Position 1 (J23101_mARi-A) | mARi expression cassette | tctggtgggtctctgtccTATAAACGCAGAAAGGCCCACCCGAAGGTGAGCCAGTGTGACTCTAGTAGAGAGCGTTCAC CGACAAAAACAGATAAAACGAAAGGCCCAGTCTTTCGACTGAGCCTTTCGTTTTATTTGATGCCTGGCTCGAGCT CGAGAAAAAAAGCCCGGACGACTGTTCGGGCTTGTCTTTTTATATGTTGGAAAATCAGTGGCAATGCAATGGCC CAACAGAAAGTTGAACACCGTCTCAGGTAAGTATCAGTTGTAAAGCTAGCATAATACCTAGGACTGAGCTAGCT GTAAAggctcgggagacctatcg | MT912081 |
| $P_{J23101}$_mARi-A Position 2 | mARi expression cassette | tctggtgggtctctgtccTATAAACGCAGAAAGGCCCACCCGAAGGTGAGCCAGTGTGACTCTAGTAGAGAGCGTTCAC CGACAAAAACAGATAAAACGAAAGGCCCAGTCTTTCGACTGAGCCTTTCGTTTTATTTGATGCCTGGCTCGAGCT CGAGAAAAAAAGCCCGGACGACTGTTCGGGCTTGTCTTTTTATATGTTGGAAAATCAGTGGCAATGCAATGGCC | MT912082 |

| | | | |
|---|---|---|---|
| | | CAACAGAAAGTATCAGTTGTAAAAAGAGGGGAAATGCTAGCATAATACCTAGGACTGAGCTAGCTGTAAAggctcgggagacctatcg | |
| $P_{J23101}$_mARi-A Position 3 | mARi expression cassette | tctggtgggtctctgtccTATAAACGCAGAAAGGCCCACCCGAAGGTGAGCCAGTGTGACTCTAGTAGAGAGCGTTCACCGACAAAAACAGATAAAACGAAAGGCCCAGTCTTTCGACTGAGCCTTTCGTTTTATTTGATGCCTGGCTCGAGCTCGAGAAAAAAAGCCCGGACGACTGTTCGGGCTTGTCTTTTTATATGTTGGAAAATCAGTGGCAATGCAATGGCCCAACAGAAAGTAAAAAGAGGGGAAATAGTCCATGGCTAGCATAATACCTAGGACTGAGCTAGCTGTAAAggctcgggagacctatcg | MT912083 |
| $P_{J23101}$_mARi-A Position 4 | mARi expression cassette | tctggtgggtctctgtccTATAAACGCAGAAAGGCCCACCCGAAGGTGAGCCAGTGTGACTCTAGTAGAGAGCGTTCACCGACAAAAACAGATAAAACGAAAGGCCCAGTCTTTCGACTGAGCCTTTCGTTTTATTTGATGCCTGGCTCGAGCTCGAGAAAAAAAGCCCGGACGACTGTTCGGGCTTGTCTTTTTATATGTTGGAAAATCAGTGGCAATGCAATGGCCCAACAGAAAATGCGTAAAGGCGAAGAACTGTTGCTAGCATAATACCTAGGACTGAGCTAGCTGTAAAggctcgggagacctatcg | MT912084 |
| $P_{J23119}$_mARi-A | mARi expression cassette | tctggtgggtctctgtccTATAAACGCAGAAAGGCCCACCCGAAGGTGAGCCAGTGTGACTCTAGTAGAGAGCGTTCACCGACAAAAACAGATAAAACGAAAGGCCCAGTCTTTCGACTGAGCCTTTCGTTTTATTTGATGCCTGGCTCGAGCTCGAGAAAAAAAGCCCGGACGACTGTTCGGGCTTGTCTTTTTATATGTTGGAAAATCAGTGGCAATGCAATGGCCCAACAGAAAGTTGAACACCGTCTCAGGTAAGTATCAGTTGTAAAGCTAGCATTATACCTAGGACTGAGCTAGCTGTCAAggctcgggagacctatcg | MT912085 |
| $P_{J23111}$_mARi-A | mARi expression cassette | tctggtgggtctctgtccTATAAACGCAGAAAGGCCCACCCGAAGGTGAGCCAGTGTGACTCTAGTAGAGAGCGTTCACCGACAAAAACAGATAAAACGAAAGGCCCAGTCTTTCGACTGAGCCTTTCGTTTTATTTGATGCCTGGCTCGAGCTCGAGAAAAAAAGCCCGGACGACTGTTCGGGCTTGTCTTTTTATATGTTGGAAAATCAGTGGCAATGCAATGGCCCAACAGAAAGTTGAACACCGTCTCAGGTAAGTATCAGTTGTAAAGCTAGCACTATACCTAGGACTGAGCTAGCCGTCAAggctcgggagacctatcg | MT912086 |
| $P_{J23104}$_mARi-A | mARi expression cassette | tctggtgggtctctgtccTATAAACGCAGAAAGGCCCACCCGAAGGTGAGCCAGTGTGACTCTAGTAGAGAGCGTTCACCGACAAAAACAGATAAAACGAAAGGCCCAGTCTTTCGACTGAGCCTTTCGTTTTATTTGATGCCTGGCTCGAGCTCGAGAAAAAAAGCCCGGACGACTGTTCGGGCTTGTCTTTTTATATGTTGGAAAATCAGTGGCAATGCAATGGCCCAACAGAAAGTTGAACACCGTCTCAGGTAAGTATCAGTTGTAAAGCTAGCACAATACCTAGGACTGAGCTAGCTGTCAAggctcgggagacctatcg | MT912087 |
| $P_{J23119}$_mARi-B | mARi expression cassette | tctggtgggtctctgtccGGACCAAAACGAAAAAACACCCTTTCGGGTGTCTTTTCTGGAATTTGGTACCGAGAAAAAAGCCCGGACGACTGTTCGGGCTTGTCTTTTTATATGTTGGAAAATCAGTGGCAATGCAATGGCCCAACAGAAAGGTATCTCGTGGTCTGACGGTAAAATCTATTGTAAGCTAGCATTATACCTAGGACTGAGCTAGCTGTCAAggctcgggagacctatcg | MT912088 |

| | | | |
|---|---|---|---|
| $P_{J23119}$_mARi-C | mARi expression cassette | tctggtgggtctctgtccTATAAACGCAGAAAGGCCCACCCGAAGGTGAGCCAGTGTGACTCTAGTAGAGAGCGTTCACCGACAAAAACAGATAAAACGAAAGGCCCAGTCTTTCGACTGAGCCTTTCGTTTTATTTGATGCCTGGCTCGAGCTCGAGAAAAAAAGCCCGGACGACTGTTCGGGCTTGTCTTTTTATATGTTGGAAAATCAGTGGCAATGCAATGGCCCAACAGAAAGTGTTACTATTGGCTGAGATAAGGGTAGCAGAAAAGCTAGCATTATACCTAGGACTGAGCTAGCTGTCAAggctcgggagacctatcg | MT912089 |
| $P_{J23119}$_mARi-D | mARi expression cassette | tctggtgggtctctgtccGGACCAAAACGAAAAAACACCCTTTCGGGTGTCTTTTCTGGAATTTGGTACCGAGAAAAAAAGCCCGGACGACTGTTCGGGCTTGTCTTTTTATATGTTGGAAAATCAGTGGCAATGCAATGGCCCAACAGAAAGTTATACCCTGTCCCGGCTGTGTATTCTATGTAAAGCTAGCATTATACCTAGGACTGAGCTAGCTGTCAAAGATTTTCCAGAGTAGGCACTTCAGGTCTATTATCACAACGAGATGGTAGACCAGAAACAAAAAAACACCCCGAAGGGTGTTCAATAATTGGggctcgggagacctatcg | MT912090 |
| $P_{J23119}$_mARi-E | mARi expression cassette | tctggtgggtctctgtccGGACCAAAACGAAAAAACACCCTTTCGGGTGTCTTTTCTGGAATTTGGTACCGAGAAAAAAAGCCCGGACGACTGTTCGGGCTTGTCTTTTTATATGTTGGAAAATCAGTGGCAATGCAATGGCCCAACAGAAAGTTATACAGCGTCTTGACGATAAAACATATGTAAAGCTAGCATTATACCTAGGACTGAGCTAGCTGTCAAAGATTTTCCAGAGTAGGCACTTCAGGTCTATTATCACAACGAGATGGTAGACCAGAAACAAAAAAACACCCCGAAGGGTGTTCAATAATTGGggctcgggagacctatcg | MT912091 |
| sfGFP_no terminator | Reporter | tctggtgggtctctgtcc**ATG**CGTAAAGGCGAAGAACTGTTCACGGGCGTAGTTCCGATTCTGGTCGAGCTGGACGGCGATGTGAACGGTCATAAGTTTAGCGTTCGCGGTGAAGGTGAGGGCGACGCGACCAACGGCAAACTGACCCTGAAGTTCATCTGCACCACCGGTAAACTGCCGGTGCCTTGGCCGACCTTGGTGACGACGTTGACGTATGGCGTGCAGTGTTTTGCGCGTTATCCGGACCACATGAAACAACACGATTTCTTCAAATCTGCGATGCCGGAGGGTTACGTCCAGGAGCGTACCATTTCCTTCAAGGATGATGGCACTTACAAAACTCGCGCAGAGGTTAAGTTTGAAGGTGACACGCTGGTCAATCGTATCGAATTGAAGGGTATCGACTTTAAAGAGGATGGTAACATTCTGGGCCATAAACTGGAGTATAACTTCAACAGCCATAATGTTTACATTACGGCAGACAAGCAAAAGAACGGCATCAAGGCCAATTTCAAGATTCGCCACAATGTTGAGGACGGTAGCGTCCAACTGGCCGACCATTACCAGCAGAACACCCCAATTGGTGACGGTCCGGTTTTGCTGCCGGATAATCACTATCTGAGCACCCAAAGCGTGCTGAGCAAAGATCCGAACGAAAAACGTGATCACATGGTCCTGCTGGAATTTGTGACCGCTGCGGGCATCACCCACGGTATGGACGAGCTGTATAAGCGTCCGTAAggctcgggagacctatcg | MT912092 |
| sfGFP_B15 | Reporter | tctggtgggtctctgtcc**ATG**CGTAAAGGCGAAGAACTGTTCACGGGCGTAGTTCCGATTCTGGTCGAGCTGGACGGCGATGTGAACGGTCATAAGTTTAGCGTTCGCGGTGAAGGTGAGGGCGACGCGACCAACGGCAAACTGACCCTGAAGTTCATCTGCACCACCGGTAAACTGCCGGTGCCTTGGCCGACCTTGGTGACGACGTTGACGTATGGCGTGCAGTGTTTTGCGCGTTATCCGGACCACATGAAACAACACGATTTCTTCAAATCTGCGATGCCGGAGGGTTACGTCCAGGAGCGTACCATTTCCTTCAAGGATGATGGCACTTACAAAACTCGCGCAGAGGTTAAGTTTGAAGGTGACACGCTGGTCAATCGTATCGAATTGAAGGGTATCGACTTTAAAGAGGATGGTAACATTCTGGGCCATAAACTGGAGTATAACTTCAACAGCCATAATGTTTACATTACGGCAGACAAGCAAAAGAACGGCATCAAGGCCAATTTCAAGATTCGCC | MT912093 |

| | | | |
|---|---|---|---|
| | | ACAATGTTGAGGACGGTAGCGTCCAACTGGCCGACCATTACCAGCAGAACACCCCAATTGGTGACGGTCCGGTT<br>TTGCTGCCGGATAATCACTATCTGAGCACCCAAAGCGTGCTGAGCAAAGATCCGAACGAAAAACGTGATCACAT<br>GGTCCTGCTGGAATTTGTGACCGCTGCGGGCATCACCCACGGTATGGACGAGCTGTATAAGCGTCCGTAATAAT<br>ACTAGAGCCAGGCATCAAATAAAACGAAAGGCTCAGTCGAAAGACTGGGCCTTTCGTTTTATCTGTTGTTTGTCG<br>GTGAACGCTCTCTACTAGAGTCACACTGGCTCACCTTCGGGTGGGCCTTTCTGCGTTTATAggctcgggagacctatcg | |
| sfGFP-<br>ECK120033737_<br>Term | Reporter | tctggtgggtctctgtcc**ATG**CGTAAAGGCGAAGAACTGTTCACGGGCGTAGTTCCGATTCTGGTCGAGCTGGACGGC<br>GATGTGAACGGTCATAAGTTTAGCGTTCGCGGTGAAGGTGAGGGCGACGCGACCAACGGCAAACTGACCCTGA<br>AGTTCATCTGCACCACCGGTAAACTGCCGGTGCCTTGGCCGACCTTGGTGACGACGTTGACGTATGGCGTGCAG<br>TGTTTTGCGCGTTATCCGGACCACATGAAACAACACGATTTCTTCAAATCTGCGATGCCGGAGGGTTACGTCCAG<br>GAGCGTACCATTTCCTTCAAGGATGATGGCACTTACAAAACTCGCGCAGAGGTTAAGTTTGAAGGTGACACGCT<br>GGTCAATCGTATCGAATTGAAGGGTATCGACTTTAAAGAGGATGGTAACATTCTGGGCCATAAACTGGAGTATA<br>ACTTCAACAGCCATAATGTTTACATTACGGCAGACAAGCAAAAGAACGGCATCAAGGCCAATTTCAAGATTCGCC<br>ACAATGTTGAGGACGGTAGCGTCCAACTGGCCGACCATTACCAGCAGAACACCCCAATTGGTGACGGTCCGGTT<br>TTGCTGCCGGATAATCACTATCTGAGCACCCAAAGCGTGCTGAGCAAAGATCCGAACGAAAAACGTGATCACAT<br>GGTCCTGCTGGAATTTGTGACCGCTGCGGGCATCACCCACGGTATGGACGAGCTGTATAAGCGTCCGTAATAAC<br>GCTGATAGTGCTAGTGTAGATCGCTACTAGAGGGAAACACAGAAAAAAGCCCGCACCTGACAGTGCGGGCTTTT<br>TTTTTCGACCAAAGGTACTggctcgggagacctatcg | MT912094 |
| mCherry-<br>B15_Term | Reporter | tctggtgggtctctgtcc**ATG**GTGAGCAAGGGCGAGGAGGATAACATGGCCATCATCAAGGAGTTCATGCGCTTCAAG<br>GTGCACATGGAGGGCTCCGTGAACGGCCACGAGTTCGAGATCGAGGGCGAGGGCGAGGGCCGCCCCTACGAG<br>GGCACCCAGACCGCCAAGCTGAAGGTGACCAAGGGTGGCCCCCTGCCCTTCGCCTGGGACATCCTGTCCCCTCA<br>GTTCATGTACGGCTCCAAGGCCTACGTGAAGCACCCCGCCGACATCCCCGACTACTTGAAGCTGTCCTTCCCCGA<br>GGGCTTCAAGTGGGAGCGCGTGATGAACTTCGAGGACGGCGGCGTGGTGACCGTGACCCAGGACTCCTCCTTG<br>CAGGACGGCGAGTTCATCTACAAGGTGAAGCTGCGCGGCACCAACTTCCCCTCCGACGGCCCCGTAATGCAGAA<br>GAAGACCATGGGCTGGGAGGCCTCCTCCGAGCGGATGTACCCCGAGGACGGCGCCCTGAAGGGCGAGATCAA<br>GCAGAGGCTGAAGCTGAAGGACGGCGGCCACTACGACGCTGAGGTCAAGACCACCTACAAGGCCAAGAAGCCC<br>GTGCAGCTGCCCGGCGCCTACAACGTCAACATCAAGTTGGACATCACCTCCCACAACGAGGACTACACCATCGTG<br>GAACAGTACGAACGCGCCGAGGGCCGCCACTCCACCGGCGGCATGGACGAGCTGTACAAGTAATAATACTAGA<br>GCCAGGCATCAAATAAAACGAAAGGCTCAGTCGAAAGACTGGGCCTTTCGTTTTATCTGTTGTTTGTCGGTGAAC<br>GCTCTCTACTAGAGTCACACTGGCTCACCTTCGGGTGGGCCTTTCTGCGTTTATAggctcgggagacctatcg | MT912095 |
| mCherry-<br>B14_Term | Reporter | tctggtgggtctctgtcc**ATG**GTGAGCAAGGGCGAGGAGGATAACATGGCCATCATCAAGGAGTTCATGCGCTTCAAG<br>GTGCACATGGAGGGCTCCGTGAACGGCCACGAGTTCGAGATCGAGGGCGAGGGCGAGGGCCGCCCCTACGAG<br>GGCACCCAGACCGCCAAGCTGAAGGTGACCAAGGGTGGCCCCCTGCCCTTCGCCTGGGACATCCTGTCCCCTCA<br>GTTCATGTACGGCTCCAAGGCCTACGTGAAGCACCCCGCCGACATCCCCGACTACTTGAAGCTGTCCTTCCCCGA | MT912096 |

| | | | |
|---|---|---|---|
| | | GGGCTTCAAGTGGGAGCGCGTGATGAACTTCGAGGACGGCGGCGTGGTGACCGTGACCCAGGACTCCTCCTTGCAGGACGGCGAGTTCATCTACAAGGTGAAGCTGCGCGGCACCAACTTCCCCTCCGACGGCCCCGTAATGCAGAAGAAGACCATGGGCTGGGAGGCCTCCTCCGAGCGGATGTACCCCGAGGACGGCGCCCTGAAGGGCGAGATCAAGCAGAGGCTGAAGCTGAAGGACGGCGGCCACTACGACGCTGAGGTCAAGACCACCTACAAGGCCAAGAAGCCCGTGCAGCTGCCCGGCGCCTACAACGTCAACATCAAGTTGGACATCACCTCCCACAACGAGGACTACACCATCGTGGAACAGTACGAACGCGCCGAGGGCCGCCACTCCACCGGCGGCATGGACGAGCTGTACAAGTAAGGCTCGATCACGGCACTACACTCGTTGCTTTATCGGTATTGTTATTACAGAGTCCTCACACTGGCTCACCTTCGGGTGGGCCTTTCTGCGTTTATATACTAGAGAGAGAATATAAAAAGCCAGATTATTAATCCGGCTTTTTTATTATTTggctcgggagacctatcg | |
| mTagBFP-B15_term | Reporter | tctggtgggtctctgtcc**ATG**TCCGAGTTGATCAAAGAGAACATGCATATGAAATTATATATGGAAGGCACTGTAGATAATCATCATTTTAAATGTACGTCGGAAGGCGAAGGTAAACCATATGAAGGTACGCAGACGATGCGCATCAAGGTGGTGGAGGGCGGTCCGCTGCCATTCGCTTTCGATATTTTAGCCACGAGCTTCCTCTACGGTTCTAAAACTTTCATCAATCACACGCAGGGTATTCCGGACTTCTTTAAACAGTCGTTCCCGGAGGGTTTCACCTGGGAACGCGTTACCACGTATGAAGATGGTGGTGTGCTTACGGCAACGCAGGACACGAGCCTTCAGGATGGGTGTTTGATTTACAACGTGAAAATTCGTGGTGTGAACTTCACGTCTAACGGCCCGGTGATGCAGAAAAAAACACTGGGTTGGGAAGCCTTTACCGAAACCCTGTATCCGGCGGACGGTGGCCTGGAAGGCCGTAATGATATGGCCTTGAAATTAGTCGGCGGTTCACACCTGATCGCGAACGCGAAAACAACCTATCGTAGTAAAAAACCAGCCAAAAACCTGAAAATGCCGggcGTCTACTACGTAGACTACCGTCTGGAGCGCattAAAGAGGCGAATAATGAAACCTATGTCGAGCAGCACGAAGTTGCGGTTGCACGCTATTGCGATCTGCCCAGCAAACTGGGCCACAAGCTTAATGGTAGCTAATAATACTAGAGCCAGGCATCAAATAAAACGAAAGGCTCAGTCGAAAGACTGGGCCTTTCGTTTTATCTGTTGTTTGTCGGTGAACGCTCTCTACTAGAGTCACACTGGCTCACCTTCGGGTGGGCCTTTCTGCGTTTATAggctcgggagacctatcg | MT912097 |

[a] Coloured, lowercase DNA sequences correspond to prefix (blue) and suffix (orange). Bold DNA sequences indicate the start codon. [b] Standardised constitutive promoters consist of a synthetic terminator (L3S3P11) (8, 9), up element, core Anderson constitutive promoter (parts.igem.org), and ribozyme insulator (RiboJ) (10). The terminator L3S3P11 used in the standardised promoter was mutated to remove the *Bsa*I site (9).

**Supplementary Table 6. List of orthogonal BASIC linker sequences used in this study.**

| BASIC linker | Type | Sequence[a] (5'→3') |
|---|---|---|
| UTR-A-RBSa | UTR-RBS linker | ggctcgttgaacaccgtctcaggtaagtatcagttgtaa**aaagaggagaaa**tagtcc |
| UTR-A-RBSb | UTR-RBS linker | ggctcgttgaacaccgtctcaggtaagtatcagttgtaa**atctaaggagg**tagtcc |
| UTR-A-RBSc | UTR-RBS linker | ggctcgttgaacaccgtctcaggtaagtatcagttgtaa**aaagaggggaaa**tagtcc |
| UTR-A-RBSd | UTR-RBS linker | ggctcgttgaacaccgtctcaggtaagtatcagttgtaa**atcccaggagg**tagtcc |
| UTR-A-RBSe | UTR-RBS linker | ggctcgttgaacaccgtctcaggtaagtatcagttgtaa**atctcgggagg**tagtcc |
| UTR-B-RBSc | UTR-RBS linker | ggctcggtatctcgtggtctgacggtaaaatctattgta**aaagaggggaaa**tagtcc |
| UTR C-RBSc | UTR-RBS linker | ggctcgtgttactattggctgagataagggtagcagaaa**aaagaggggaaa**tagtcc |
| UTR D-RBSc | UTR-RBS linker | ggctcgttataccctgtcccggctgtgtattctatgtaaa**aagaggggaaa**tagtcc |
| UTR E-RBSc | UTR-RBS linker | ggctcgttatacagcgtcttgacgataaaacatatgtaa**aaagaggggaaa**tagtcc |
| MLA | Methylated linker | ggctcgggtaagaactcgcacttcgtggaaacactattatctggtgggtctctgtcc |
| MLB | Methylated linker | ggctcgggagacctatcggtaataacagtccaatctggtgtaacttcggaatcgtcc |
| L1 | Neutral linker | ggctcgttacttacgacactccgagacagtcagagggtatttattgaactagtcc |
| L2 | Neutral linker | ggctcgatcggtgtgaaaagtcagtatccagtcgtgtagttcttattacctgtcc |
| L3 | Neutral linker | ggctcgatcacggcactacactcgttgctttatcggtattgttattacagagtcc |
| L4 | Neutral linker | ggctcgacccacgactattgactgctctgagaaagttgattgttacgattagtcc |
| L5 | Neutral linker | ggctcgagaagtagtgccacagacagtattgcttacgagttgatttatcctgtcc |

[a] DNA sequence colours correspond to upstream scar (blue) and downstream scar (orange). The DNA sequence in underlined-bold indicates the RBS sequence.
